# Context-dependent abdomen bobbing in a jumping spider: a dynamic visual signal

**DOI:** 10.1101/2025.10.01.679781

**Authors:** Nadja Geiger, Chiara Hirschkorn, Marie E. Herberstein, Mikkel Roald-Arbøl, Daniela C. Rößler

## Abstract

Anti-predatory visual signals play a key role in deterring predatory attacks, and can include crypsis, warning colors or visual mimicry. Static traits such as shape or color are well-studied, whereas locomotion-related signals remain less understood despite being particularly intriguing as they can be controlled by the animal’s perception of danger. In this study, we explore the potential function of a highly conspicuous dynamic behavior—abdomen bobbing—exhibited by the diurnal jumping spider *Heliophanus* cf. *cupreus* during locomotion. Due to the high conspicuousness, we hypothesized that abdomen bobbing could function as a visual signal, specifically as an anti-predatory signal. To test this hypothesis, we first analyzed the behavior and found that bobs occur almost exclusively during gait stops (98.5%) with 83.5% of gait stops entailing abdomen bobbing. We then examined its context-dependency, measuring the prevalence of abdomen bobbing under four conditions: 1) control, 2) in darkness, 3) during prey encounters, 4) during simulated conspecific encounters (i.e. mirror), and 5) during predator encounters. Abdomen bobbing was virtually absent in darkness and during prey encounters, but significantly increased during conspecific and predator encounters, suggesting active control. The increase in abdomen bobbing when confronted with a predator strongly supports an anti-predatory signaling function, potentially a form of locomotion mimicry. We equally found support for an intra-specific signaling function, which might indicate the co-option of a signal from one context (predatory) to another (conspecific). Our study opens new avenues for investigating the role of motion in both inter- and intraspecific signaling.

## Introduction

Animals need to move in search of food, shelter or mates. However, locomotion can break an animal’s camouflage or crypsis and increase detection by predators (Tan et al., 2024). While less well understood than static and morphological anti-predatory traits, animals can counteract the increased visibility by adjusting locomotion and even using motion to conceal, dazzle and deceive predators (see Tan et al., 2024). For example, praying mantids (Watanabe & Yano, 2009) and stick insects (Bian et al., 2016) sway their bodies in an apparent mimicry of trembling leaves. By increasing the swaying in wind conditions their detection by predators may be lowered.

Locomotion can also deter predation as an extension of Batesian or Mullerian mimicry, which classically is based on static morphological resemblance of the mimic to the toxic or unpalatable model. In locomotor mimicry, the resemblance extends to how the model moves (Huang & Caro, 2023) with evidence from mimetic butterflies (e.g. Kitamura & Imafuku, 2010), wasp-mimicking clear-winged moths (Skowron Volponi et al., 2018) and wasp-mimicking hover flies (Penney et al., 2014). Complex behavioral mimicry has also been reported in ant-mimicking spiders that in addition to morphological ant resemblance (myrmecomorphy), also mimic ant behavior. For example, myrmecomorphic spiders move at speeds matching those of ants (Subramaniam et al., 2022), move in typical ant-like stop-go fashion and winding trajectories, with their first legs raised in resemblance of antennae when stopping (Shamble et al., 2017). What is particularly compelling about mimetic behavior in myrmecomorphs is the ability to modulate the behavior in different contexts. For example, *Myrmarachne* ant-mimics increase behavioral mimicry of ants when moving more and thus be potentially more exposed to predators, but abandon ant-like behavior when confronted with immediate danger (Ceccarelli, 2008). It would also be of advantage to reduce behavioral ant mimicry when in proximity of ant specialists (e.g. *Servaea* jumping spiders), as these readily attack ant mimics at similar rates to ants (Pekár et al., 2017).

A particularly intriguing behavior that has been described in myrmecomorphic spiders, and also in non-mimicking spiders (see Table 1) is abdomen bobbing (or twitching) where the spider lifts and lowers the abdomen as it walks. In particular, abdomen bobbing in myrmecomorphic spiders has been linked to how some ants also lift and lower their gaster during locomotion (Ceccarelli, 2008). Explanations for this highly conspicuous and possibly energetic behavior ranges from non-signaling physiological functions, such as aiding ventilation to signaling functions such as threat displays between males, or courtship display between males and females. Finally, abdomen bobbing in spiders may be a form of behavioral mimicry of ants and wasps functioning as anti-predatory signaling (Table 1).

**Table 1.** Collection of species with reported abdomen bobbing. Descriptions, terminology, hypotheses and references for previously investigated species.

| Species | Tribe | Term used | Context | Hypothesis | Reference |
| --- | --- | --- | --- | --- | --- |
| *Cosmophasis micarioides* | Chrysillini | abdomen bobbing<br>abdomen twitching<br>raised abdomen | normal locomotion<br>male-female interaction<br>male-male interaction | ventilation<br>courtship<br>threat display | Jackson, 1986b |
| *Cosmophasis umbratica* | Chrysillini | abdominal bobbing<br>abdomen flexed | normal locomotion<br>male-female interaction<br>male-male interaction | courtship<br>male-male display | Lim & Lee, 2004 |
| *Cyllobelus rufopictus*<br>(syn. *Natta horizontalis*) | Chrysillini | abdomen bob<br>abdomen twitch<br>raised abdomen | normal locomotion<br>male-male interaction | courtship<br>male-male display | Jackson, 1986c |
| *Siler semiglaucus* | Chrysillini | abdomen bobbing | hunting<br>feeding | active situational<br>presence | Hawes, 2018 |
| *Habronattus agilis* | Dendryphantini | abdomen twitching | courtship | vibration<br>courtship | Maddison & Stratton, 1988 |
| *Maratus speciosus* | Euophryini | opisthosomal bobbing<br>bobbing of the fan | courtship | vibration<br>courtship | Hill & Otto, 2014 |
| *Myrmarachne exasperans* | Myrmarachnini | opisthosoma movement<br>bobbing | normal locomotion | parasitoid wasp mimicry | Hurni-Cranston & Hill, 2019 |
| *Myrmarachne lupata* | Myrmarachnini | abdomen twitch<br>abdomen flutter<br>abdomen waving<br>abdomen bob | normal locomotion<br>male-female interaction | courtship<br>threat display<br>ant mimicry | Jackson, 1982, 1986a |
| *Peckhamia picata* | Sarindini | abdomen bobbing<br>abdomen lifting and<br>retreating | normal locomotion<br>predator encounter | ant-like behavior<br>defensive behavior | Durkee et al., 2011 |
| *Cyrba algerina* | Spartaeini | abdomen bob<br>abdomen twitch | normal locomotion<br>male-female interaction<br>male-male interaction | activity<br>ventilation<br>visual display | Jackson & Hallas, 1986 |
| *Brettus adonis* | Spartaeini | abdomen twitch | hunting (web invasion) | - | Jackson & Hallas, 1986 |
| *Brettus cingulatus* | Spartaeini | abdomen twitch | hunting (web invasion) | - | Jackson & Hallas, 1986 |

Here, we want to experimentally distinguish between the proposed, non-mutually exclusive functions of abdomen bobbing in the non-myrmecomorphic jumping spider, *Heliophanus* cf. *cupreus* (Supplemental Video S2). First, we describe the gait of *Heliophanus* cf. *cupreus* and compare it to that of two other common salticids (*Salticus scenicus* and *Marpissa muscosa*). We then experimentally manipulate the context in which the spider moves. We first determine whether abdomen bobbing has a non-signaling or a signaling function. If abdomen bobbing has a signaling function, we predict that this diurnal spider will engage in abdomen bobbing in the light, while it should cease to do so in the dark. In further experiments we test the prediction that abdomen bobbing has a visual function such as an anti-predatory signal. In this case, we predict that bobbing increases when exposed to a predator, while at the same time we expect a decrease of the behavior in a foraging context, as being conspicuous during stalking of prey could hamper prey capture. Lastly, we test for an intra-specific signaling function for which we predict that abdomen bobbing would increase in the presence of a conspecific.

## Material and methods

### Study species and animal husbandry

*Heliophanus* cf. *cupreus* (Walckenaer & Fabricius, 1802) (hereafter *Heliophanus*) is a species of jumping spider distributed across Europe, Northern Africa, and parts of Asia and can usually be found on low vegetation (World Spider Catalog, 2025). The opisthosoma is dark with bright spots, covered in iridescent hairs. The adult female measures about 4.6-5.8 mm in body length with bright yellow pedipalps in contrast to their darker body coloration. Adult males are smaller, ranging from 3.6-4.0 mm in body length, and have dark-colored pedipalps (Bellmann, 2016; Metzner, 2024; Wesołowska, 1986).

All individuals used in the experiments were wild-caught animals collected in Constance, Germany (47° 41’ 28.2264” N 9° 11’ 10.3164” E) between 2021 and 2023. The spiders were transported to the laboratory and kept separately in clear plastic boxes (5.5 × 5.5 × 8.5 cm, Amac). Each box was enriched with a plywood board (5.4 × 5.4 cm) to provide structure and shelter. The animals were fed with flightless *Drosophila melanogaster* twice a week *ad libitum* and had access to water provided in an Eppendorf tube (2ml) filled with water and stoppered with a cotton ball. Animals were housed under a constant temperature of 22°C (+/-2 degrees Celsius) and under a 12:12 light:dark cycle.

### Experimental design

A total of 102 spiders (juveniles = 31, females = 52, males = 29) were placed on a 3D-printed horizontal runway and subsequently filmed moving along the runway. Runways were mounted on a 12.8 cm pole in front of a camera and the length and width of the runway was adjusted for each treatment to optimize stimulus encounter (see Fig. 1). In all trials the runway was covered with white filter paper, which was removed and the runway cleaned with ethanol (98%) after each trial to remove any potential silk and/or chemical traces left. To maximize sample size in each treatment, some individuals were reused, resulting in repeated measures, which was accounted for in the statistical analyses (see *Video scoring and data analysis* below).

**Figure 1.**
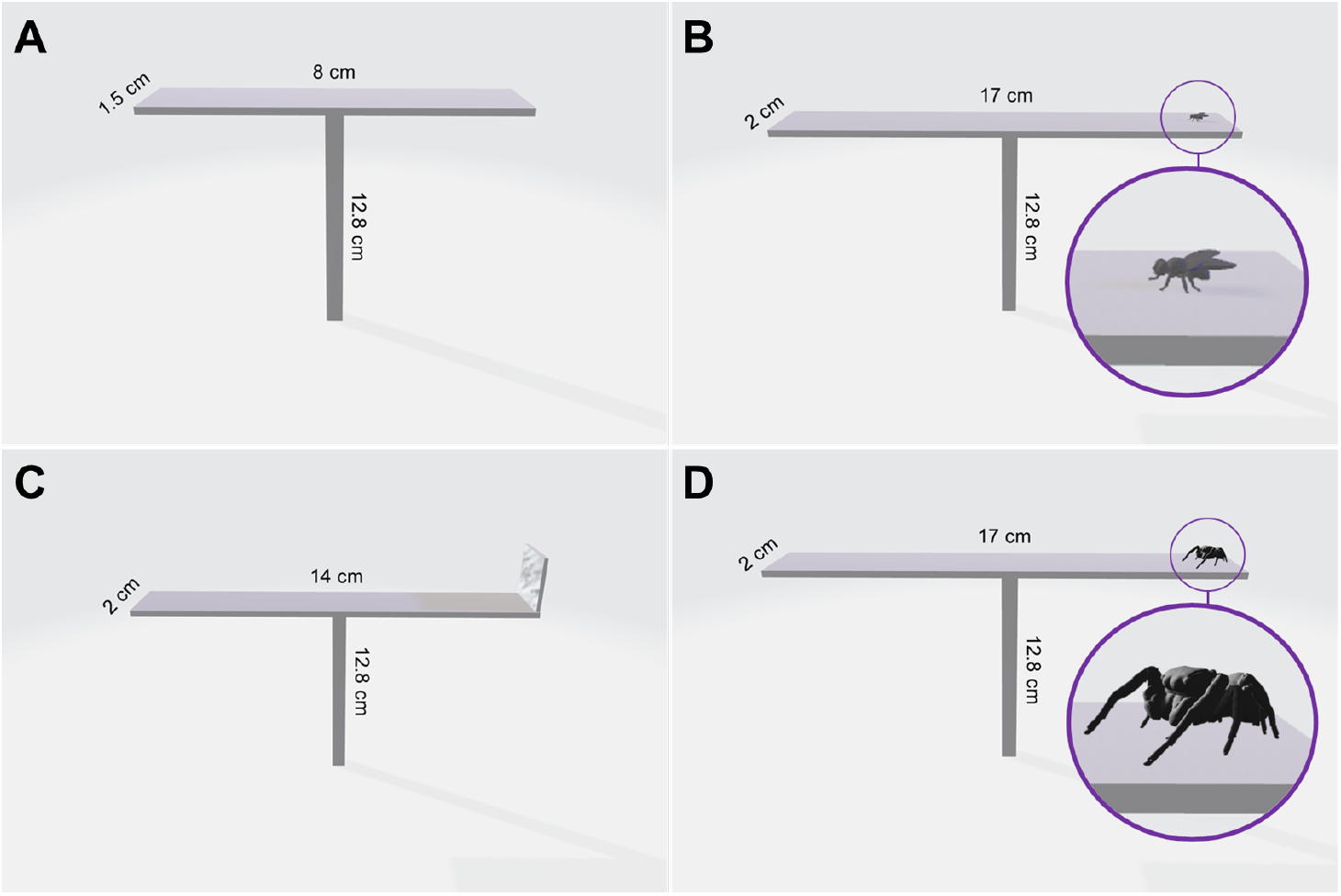
Experimental setups. Treatments: A) Control and dark, B) foraging, C) conspecific (i.e., mirror), D) predator (3D model). Illustration: Daniela C. Rößler.

All trials under daylight were filmed with a Nikon D7500 (Nikon, Tokyo, Japan) under LED light strips (daytime eco110.2 Ultra White 7.000 K). For trials run in darkness, we used infrared lights (INSTAR IN-907) and a full-spectrum modified DSLR camera (Nikon D7200, Tokyo, Japan) for infrared video recording.

During the control and light context trials the scoring started once the spider entered the runway and lasted a maximum of three minutes. Spiders were manually transferred from a plastic vial using a piece of paper or wood and encouraged to jump onto the runway by holding the paper close to it. If this failed, spiders were gently tapped out of the vial from above and placed onto the runway. A trial ended earlier when the spiders jumped off. For the foraging-, conspecific-, or predator trials, the scoring started once the spider visually encountered the presented stimulus (fly, mirror, or 3D model of a predator) at the end of the runway and ended once the spider attacked (fly), retreated, jumped off or after three minutes passed.

### Control

We quantified a baseline (control) of bobbing behavior of *Heliophanus* during daylight conditions and in the absence of a stimulus, testing 55 spiders (juveniles = 12, females = 27, males = 16) in a randomized order on a 8cm × 1.5 cm runway platform (Fig. 1A). We additionally filmed two other jumping spider species: *Marpissa muscosa* and *Salticus scenicus* as species-controls using the same experimental setup. We tested 10 female *Marpissa muscosa* and 17 female *Salticus scenicus*.

### Context-dependent tests

We tested *Heliophanus* under different treatments and compared abdomen bobbing to the control trials (Supplemental Video S1).

### Light context (darkness)

To test if abdomen bobbing represents a visual signal, we tested whether bobbing behavior persists in the absence of visible light, using the same setup as the control treatment, but with infrared lights (Fig. 1A). A total number of 35 spiders (juveniles = 6, females = 16, males = 13) were chosen at random and tested in a randomized order.

### Foraging context

To test whether the abdomen bobbing might function as a signal in prey capture, we observed spiders in the presence of tethered flightless *Drosophila melanogaster*. For each trial, a live fly was fixed to one end of the runway (Fig 1B) using double sided sticky tape which held the flies in place while allowing them to move as they tried to free themselves. The movement reliably triggered stalking and ultimately an attack response in the spiders. The spider was placed at one end of the runway with the fly at the opposite end. The trial commenced when the fly was visually detected by the spider, indicated by a freeze behavior, while eyes were directed at the fly. A total of 25 spiders (juveniles = 4, females = 11, males = 10) were chosen at random and tested in randomized order.

### Conspecific context

To test whether abdomen bobbing behavior functions as an intra-specific signal, we presented the test spider with a mirror (2 × 2 cm) at the end of the runway (Fig. 1C). Once the spider detected its mirror image, it would move towards it. A total of 9 spiders (juveniles = 5, female = 4) were randomly chosen and tested in randomized order.

### Predator context

To test whether abdomen bobbing functions as an anti-predatory signal, we exposed test spiders to 3D-printed models (11 mm in length) of a larger predatory jumping spider (3D prints based on microCT scans of *Phidippus audax*, stl-file available from Zenodo: Rößler & Shamble, 2022) (Plate & Rößler, 2024; Rößler et al., 2022). The eyes of the models were painted with black acrylic paint (Revell). The models were placed at one end of the runway (Fig. 1D) and the test spider released at the opposite end. A total of 18 spiders (juveniles = 8, females = 9, males = 1) were randomly chosen and tested in randomized order.

### Video tracking

In order to illustrate the movement of the abdomen during a walking sequence (Fig. 2), we used the pose estimation software SLEAP (Pereira et al., 2022) to track spinnerets and the anterior-superior edge of the abdomen. Trajectories were processed with the animovement R package (Roald-Arbøl, 2024), and spiders were determined to be walking when moving at > 20 mm/s.

**Figure 2.**
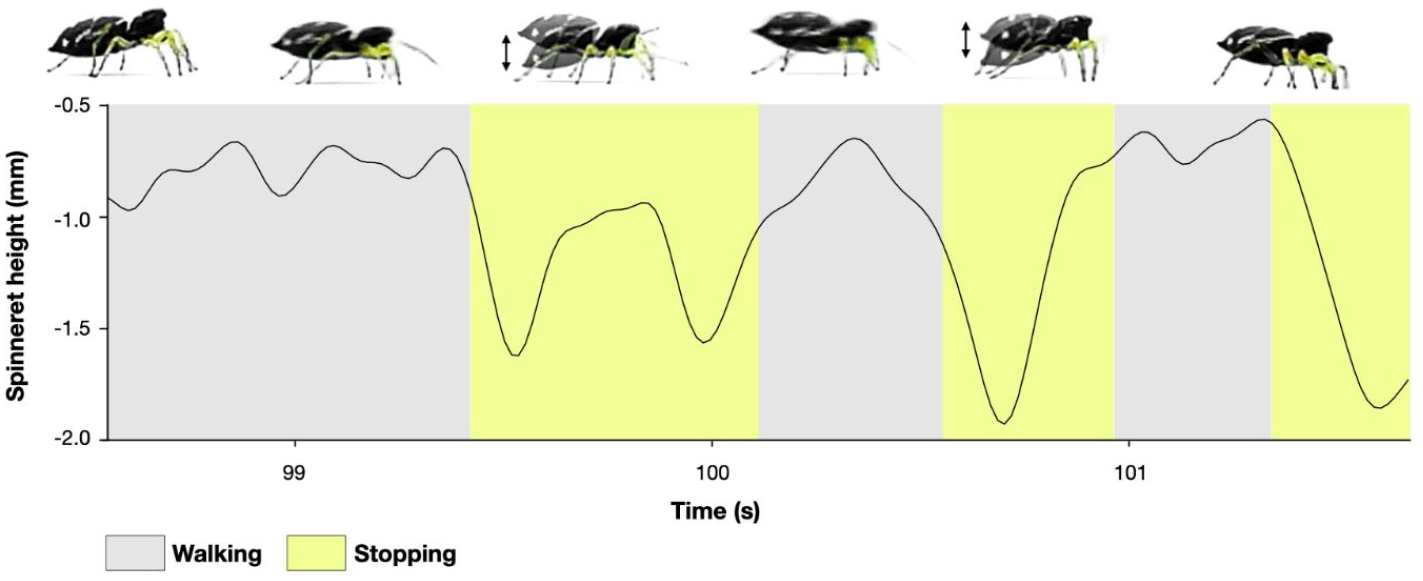
Behavioral sequence. The graph shows the vertical distance of the spinnerets relative to the most anterior-superior end of the abdomen over time (tracked using SLEAP). Colors indicate whether the spider was walking or stopping. A spider was considered to be walking if it moved at > 20 mm/s. Sequence of frames on top show *Heliophanus* during walking in the control trial (no stimulus). Each frame captures key moments in the sequence, illustrating the down and up motion during gait stops.

### Video scoring and data analysis

All videos were manually scored using BORIS (Friard & Gamba, 2016) according to the observed behaviors defined in Table 2.1. The movement of the abdomen was measured by recording each time the abdomen reached its highest and/or lowest position, as well as whether the spider was walking or had stopped.

**Table 2.** Behavioral ethogram. Descriptions and definitions for manually scored behaviors.

| Scored Behavior | Behavior Type | Description |
| --- | --- | --- |
| Walking | State Event | The spider is walking on the runway |
| Stopping | State Event | The spider is not walking (appendages may still move, but the spider's overall velocity is zero) |
| Abdomen up | Point Event | The spider's abdomen reaches the highest point in an upward movement |
| Abdomen down | Point Event | The spider's abdomen reaches the lowest point in a downward movement |
| Silk Anchoring | Point Event | The spider is casting a silk anchor on the runway |
| Stimulus detection | Point Event | The stimulus is in the sightline of the spider. This behavior marks the start of the trial in treatments with a stimulus |

To account for the difference of trial length, we randomly selected three stops for each trial, from which abdomen bobs per stop were calculated. For trials with less than three stops we used the maximum number of stops for that trial. Stops that lasted longer than 15 seconds were not included in the analysis, as we could not reliably interpret these as gait stops.

Statistical analyses were carried out in R 4.3.1 (R Core Team, 2024). We used generalized linear mixed models (GLMM) with the package *glmmTMB* (Brooks et al., 2017) to test whether treatments had significant effects on the dependent variables (abdomen bobbing or stop duration). We then applied an analysis of deviance to the resulting models using the package *car* (Fox & Weisberg, 2019). Subject ID was always included as a random factor to account for multiple data points from each individual as well as for repeated trials for some individuals. Post hoc analyses with Bonferroni correction were conducted with the package *emmeans* (Lenth, 2024). Model fit was confirmed using the package *DHARMa* (Hartig, 2022). Unless stated otherwise Gamma loglink was the best fit for models using duration, when dealing with count data (abdomen bobs) we used a poisson distribution. Plots were generated using the package *ggplot2* (Wickham, 2016). All values are median ± IQR unless specified otherwise. The R script for analyses of manually scored data along with experimental data are available in the supplements (Supplemental Data S1).

### Ethics

All experiments were conducted on invertebrates for which ethics approval is not required. We adhered to the guidelines for the treatment of animals in behavioral research by the Association for the Study of Animal Behaviour (ASAB) (Buchanan et al., 2012).

## Results

### Description of abdomen bobbing

The walking bouts during control trials shown by *Heliophanus* lasted for 1.08 ± 1.87 s (n_obs_ = 539) and were interrupted by gait stops (stops < 15 s), lasting on average 0.67 ± 1.4 s (median and IQR) (n_obs_ = 466). Abdomen bobbing was regularly performed during normal walking. The bobbing movement consisted of a lowering and lifting of the abdomen. In the control trials, abdomen bobs were observed in 83.5% of gait stops, occurring either once or multiple times, with an average of 1.85 ± 1.89 (mean ± SD) bobs per stop (n_obs_ = 389).

To demonstrate that bobbing was not a general salticid trait, we tested individuals of *Marpissa muscosa* (n = 10) and *Salticus scenicus* (n = 17) on the same setup, comparing them to the *Heliophanus* control. In both species, abdominal bobbing was completely absent allowing no statistical comparison.

However, there was a significant difference of stop duration between the three tested spider species (Fig. 3, GLMM Analysis of deviance, χ2 = 83.48, df = 2, **p < 0.0001**, n_obs_ = 571; n_ind_ = 78). *Heliophanus* evidently had the shortest stop duration (median ± IQR; 0.67 ± 1.4 s, n_obs_ = 466), while *Salticus* (4.32 ± 6.42 s, n_obs_ = 82) and *Marpissa* (2.74 ± 4.89 s, n_obs_ = 23) had significantly longer stops (Post hoc, *Heliophanus*/*Salticus*: ratio = 0.21, SE = 0.04, **p < 0.0001**; *Heliphanus*/*Marpissa*: ratio = 0.3, SE = 0.09, **p = 0.0001**). There was no significant difference in the stop duration between *Salticus* and *Marpissa* (Post hoc test, *Salticus*/*Marpissa*: ratio = 0.71, SE = 0.23, p = 0.84).

**Figure 3.**
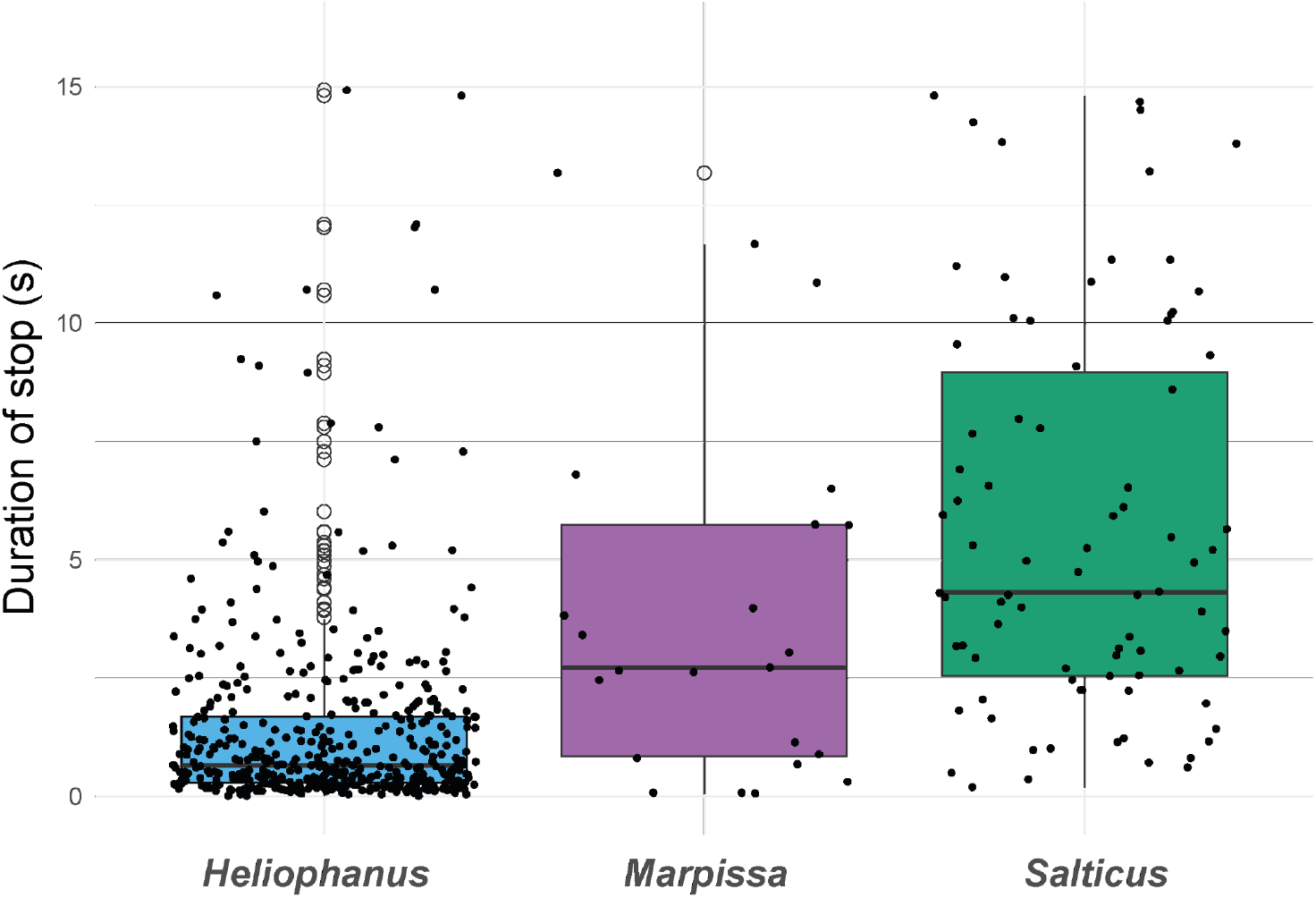
Stopping duration across three species. Boxplots showing duration of gait stops (stops lasting less than 15 seconds) for three tested spider species (*Heliophanus* cf. *cupreus*: n_obs_ = 466; n_ind_ = 55; *Salticus scenicus*: n_obs_ = 82; n_ind_ = 17; *Marpissa muscosa*: n_obs_ = 23; n_ind_ = 10). Black horizontal lines represent the median, lower and upper bound of the boxes show 25th and 75th percentiles with whiskers representing ±1.5 interquartile range. Empty circles show outliers, small dots represent single stops.

Within the control treatment of *Heliophanus*, there was no significant difference of bobbing rate per stop between males, females and juveniles (Fig. 4, GLMM Analysis of deviance, χ2 = 4.02, df = 2, p = 0.134, n = 55).

**Figure 4.**
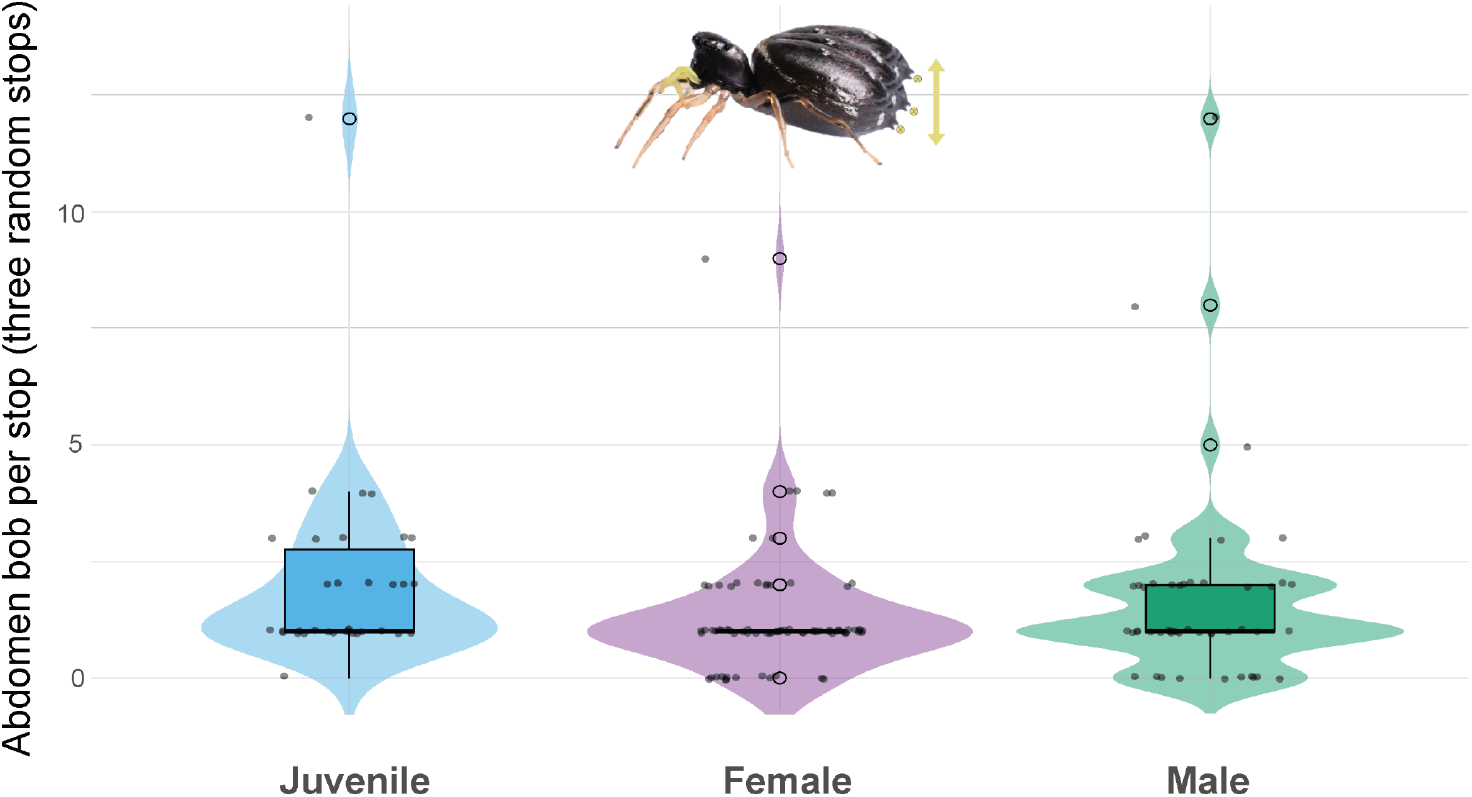
Abdomen bobbing across sexes. Boxplots showing abdomen bobs per stop for different sexes/maturity levels of *Heliophanus* in the control treatment for three random stops in each trial. Black horizontal lines represent the median, lower and upper bound of the boxes show 25th and 75th percentiles with whiskers represent ±1.5 interquartile range. Small dots represent single stops, empty circles represent outliers.

### Context-dependent abdomen bobbing

We found a significant effect of the treatment on abdomen bobbing behavior (Fig. 5, GLMM Analysis of deviance, χ2 = 184.2, df = 4, **p < 0.0001**, n_obs_ = 422; n_ind_= 102). Abdomen bobbing was virtually absent (median = 0) in the dark treatment and significantly reduced compared to the control (Post hoc, ratio = 5.62, SE = 1.24, **p < 0.0001**). Likewise, bobbing was significantly reduced in the prey capture treatment (median = 0) compared to the control (Post hoc, ratio = 7.36, SE = 2.16, **p < 0.0001**). When spiders faced a size- and sex-matched conspecific (i.e. a mirror), abdomen bobbing significantly increased compared to the control (Post hoc, ratio = 1.95, SE = 0.31, **p = 0.0004**). Bobbing also significantly increased when faced with a 3D predator model compared to the control (Post hoc, ratio = 0.34, SE = 0.05, **p < 0.0001**).

**Figure 5.**
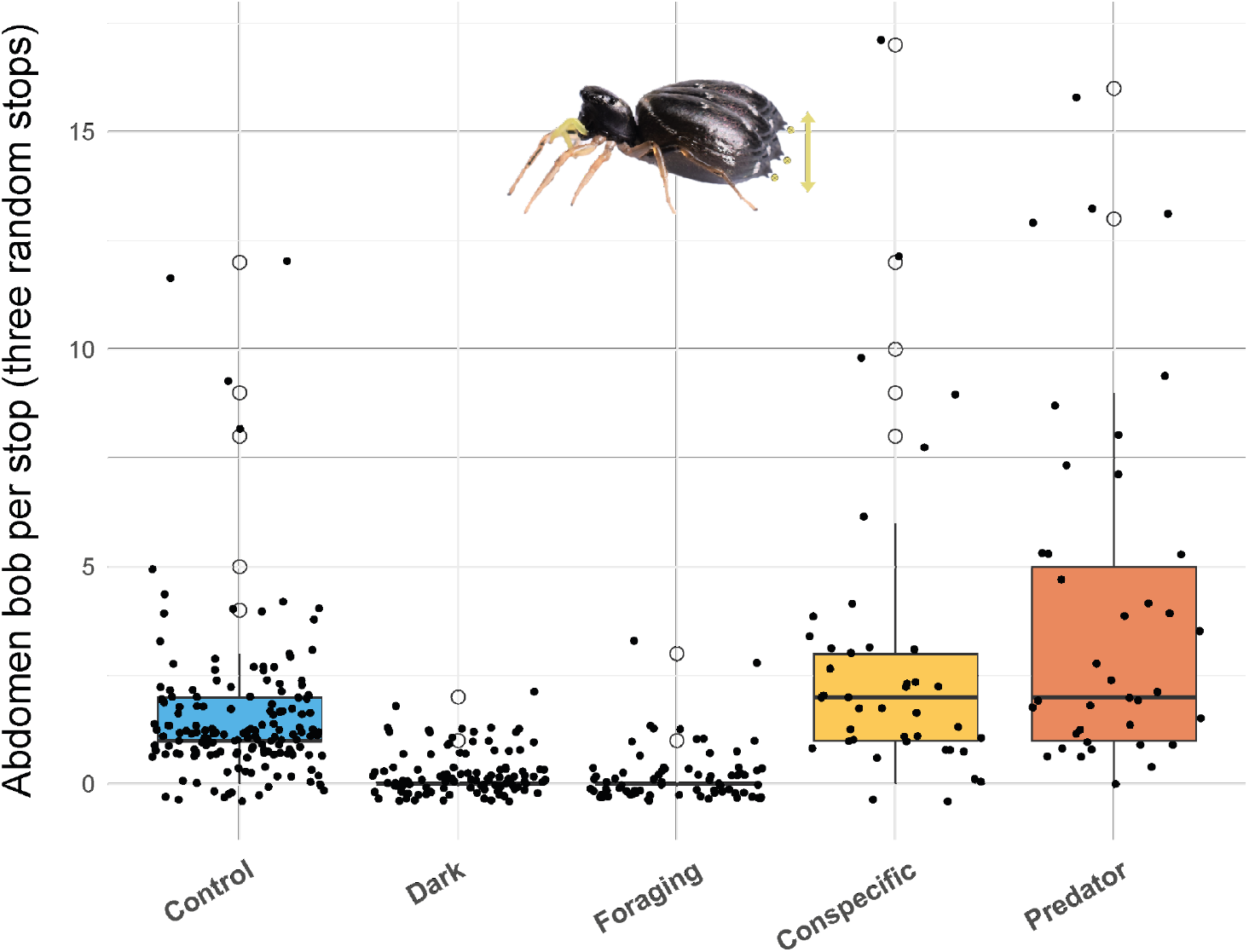
Abdomen bobbing across treatments. Boxplots showing number of abdomen bobs per stop depending on treatment. For each trial, three stops were randomly picked. Black horizontal lines represent the median, lower and upper bound of the boxes show 25th and 75th percentiles with whiskers representing ±1.5 interquartile range. Empty circles show outliers, smaller dots represent all data points.

Additionally, we investigated whether the stop duration differed between treatments (Fig. 6). There was an overall significant effect of treatment on the stop duration (GLMM Analysis of deviance, χ2 = 19.094, df = 4, **p = 0.0007**, n_obs_ = 422; n_ind_ = 102). After Bonferroni correction, the only significant difference was between the control and the predator treatment (Post hoc, Control/Predator: ratio = 0.42, SE = 0.09, p = **0.0011**) and the dark and the predator treatment (Post hoc, bonferroni-corrected, Dark/Predator: ratio = 0.41, SE = 0.1, p = **0.0014**) with stop duration being longer in the predator treatment. No other pairwise comparisons were statistically significant.

**Figure 6.**
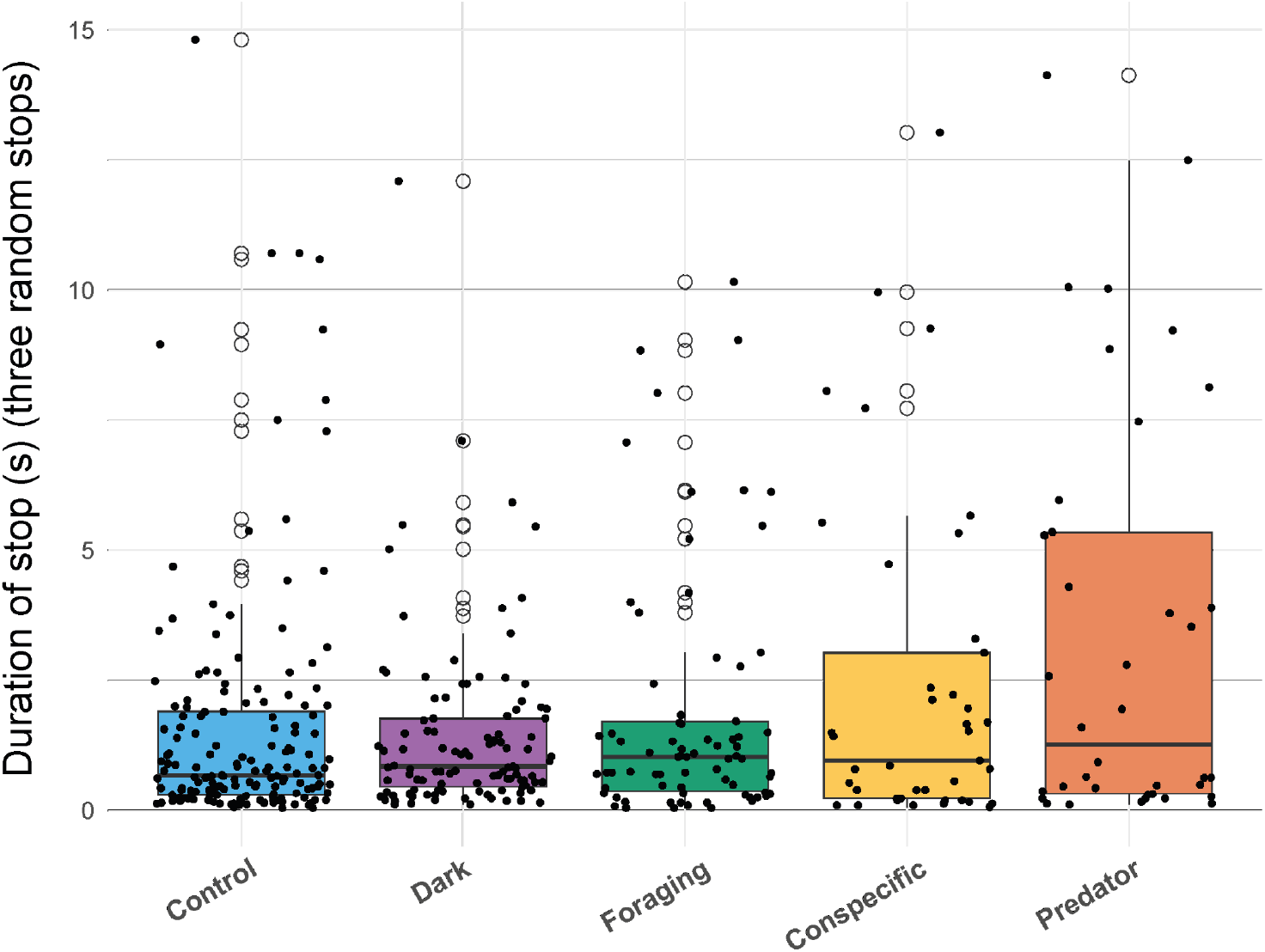
Stop duration across treatments. Boxplots showing duration of gait stops (in seconds) for each treatment. From each trial, three stops were randomly picked to account for varying trial lengths. Black horizontal lines represent the median, lower and upper bound of the boxes show 25th and 75th percentiles with whiskers represent ±1.5 interquartile range. Empty circles show outliers, smaller dots represent all data points.

## Discussion

Abdomen bobbing has been reported in several salticid species (see Table 1). However, no obvious function for abdomen bobbing has yet been determined. No systematic or ecological pattern emerges by comparing traits of salticids that exhibit abdomen bobbing: Some are ant-mimics (e.g. *Myrmarachne, Peckhamia*), others are araneophagic (e.g. *Cyrba, Brettus*), some are cryptically colored (*Brettus*) and others highly conspicuous (e.g. *Maratus*). Here we wanted to test the hypothesis that abdomen bobbing may serve as a visual signal, and who the potential recipients for such a visual signal might be.

During normal locomotion, *Heliophanus* cf. *cupreus* showed a rapid stop-and-go gait that has been described for other salticids (Jackson, 1986b, 1986c; Jackson & Hallas, 1986), however in *Heliophanus* bobs almost exclusively occurred during gait stops (98.5%), contrasting for example with *Cosmophasis micarioides* (Jackson, 1986b), where bobbing continued when the spider resumed walking (Fig. 2). Additionally, in contrast to *C. micarioides*, bobbing in *Heliophanus* generally started with a downward motion, rather than upward. While the downward motion brought the abdomen close to the ground, it never touched it and it did not involve casting of silk anchors, which were recorded as distinct events (see Table 2).

Abdomen bobbing was clearly under voluntary control with individuals increasing and decreasing it under different contexts to the point of seizing bobbing entirely. These results exclude ventilation as an explanation for the behavior as proposed in earlier studies (Jackson & Hallas, 1986). The experimental contexts we tested were designed to distinguish between several functions proposed for abdomen bobbing, generating an overwhelming support for a visual function: bobbing was entirely absent under dark, a non-visual condition. During the predator encounters however, bobbing was present, and even significantly increased compared to the control. Jumping spiders readily recognize and respond to models of larger predatory spiders, which often trigger a freeze response (see also Plate & Rößler, 2024; Rößler et al., 2022), or in the case of *Heliophanus*, an abdomen bobbing response.

Our predator-context experiment supports a possible anti-predator signal function for the abdominal bobbing in *Heliophanus*. While we did not test direct fitness impacts of bobbing behavior, similar behaviors observed in other invertebrates may support a more general anti-predator function. Male stalk-eyed flies (*Teleopsis dalmanni*) increase their chances of survival by abdomen bobbing (Worthington & Swallow, 2010) and even *Drosophila* has been reported to display abdomen lifts in response to the presence of a salticid predator (Parigi et al., 2019), though survival impact was not quantified in this studied. The Asian honeybee (*Apis cerana*) uses abdomen wagging as a form of an “I See You Signal”, deterring predator attacks (Tan et al., 2012).

The protective mechanisms may indeed be behavioral mimicry of a defended model, such as wasps and ants. Body oscillations, including abdominal wagging, have been reported for various wasp species (Savoyard et al., 1998) and abdominal wagging in fire ants (Cassill et al., 2016). While *Heliophanus* is not considered a morphological ant mimic, the very pointy tip of the abdomen (= stinger?) together with the overall glossiness of the spider’s exterior is reminiscent of the glossy, smooth abdomen that characterizes many ant-mimicking spiders (Cushing, 1997; Nelson, 2010).

Bobbing was also significantly increased in the conspecific contexts, although sample size for this experiment was somewhat low. We chose a mirror to create size- and sex-matched conspecific encounters, a method frequently used in jumping spiders (e.g. Cross et al., 2007) who readily interact with their mirror image. Conspicuous inter-specific signals, such as aposematic signals are often co-opted into an intra-specific sexual signaling context where they may indicate male or female quality (see Rojas et al., 2018). Thus, abdomen bobbing in *Heliophanus* may contain relevant information about individual quality that may determine the outcomes of intra-specific interactions, an intriguing prospect for experimental exploration.

The only visual context that reduced bobbing was the prey encounter. Once the prey item was detected, the spiders crouched down to the ground, snuck up to the prey, and attacked. It is likely that conspicuously bobbing would otherwise alert the prey to the stalking spider, hampering successful prey capture.

Our experimental work not only provides strong support for abdomen bobbing as an anti-predator signal, but also generates further hypotheses that can be tested experimentally in the lab as well as the field by quantifying the bobbing rate in different behavioral contexts, interactions and across different microhabitats. For example, bobbing could be increased in more exposed sites (i.e. on top of vegetation) and decreased when the spider is obscured by dense vegetation. Testing newly emerged spiderlings could yield insight into whether abdomen bobbing is an innate or learned behavior and testing opposite-sex encounters would help untangle the function of abdomen bobbing in interspecific communication and courtship behavior. Ultimately, testing how bobbing directly impacts fitness will add an important layer to a comprehensive understanding of this intriguing dynamic signal. Finally, a broader comparative analysis of bobbing behavior among salticid, and perhaps even other spider families, may help understand interspecific differences and identify ecological correlates for this intriguing trait

## Supporting information

RMarkdown Script

## Acknowledgements

We would like to thank Alex Jordan and Meg Crofoot for providing resources and space for us to conduct this research as well as the Zukunftskolleg for supporting us to present early parts of this research at a conference. We also thank Kaz Uyehara for support with the initial data exploration.

## Inclusion and diversity statement

The authors greatly value equity, diversity and inclusion in science (Rößler et al., 2020; Sweet, 2021). The authors represent several career stages from postgraduate to full professor. The authors come from Germany, Austria and Denmark. One or more of the authors self-identifies as a member of the LGBTQ+ community. We actively worked to promote gender balance (of first author) in our reference list.

## Conflict of interest

The authors declare no conflict of interest.

## Ethics approval

Not applicable.

## Data availability statement

All data and code are available in the online supplementary material.

## Author contributions

D.C.R. initiated the idea for the project. C.H. collected the data. D.C.R., N.G. and M.R.-A. analyzed the data and created the figures and supplemental videos. D.C.R., N.G. and M.H. wrote the manuscript with contributions from M.R.-A..

## Notes

### Competing Interest Statement

The authors have declared no competing interest.

