## Supplementary material for "Context-dependent abdomen bobbing in a jumping spider: a dynamic visual signal": RMarkdown Script

### Abdomen Bobbing

Daniela Rößler

2024-09-30

#### Libraries

```
library(ggplot2) #plots
library(DHARMa)#model fit
library(car) #anova
library(emmeans) #estimated marginal means
library(glmmTMB) #glmms
library(dplyr) #data management
library(psych) #descriptivestats
```

#### Custom colorblind friendly palette

```
##Color Palette for plots
cbbPalette <- c("#E69F00", "#56B4E9", "#009E73", "#F0E442", "#0072B2", "#D55E00", "#CC79A7")
```

#### Loading the data

```
bob <- read.csv("C:/Users/danie/Desktop/AbdomenBobbingPaper_Ethology/Supplemental Data S1/Dataset1_Thre
bobraw <- read.csv("C:/Users/danie/Desktop/AbdomenBobbingPaper_Ethology/Supplemental Data S1/Dataset2_Al
```

```
bob$Test[bob$Treatment == "Light"] <- "Control"
bob$Test[bob$Treatment == "Dark"] <- "Light Context"
bob$Test[bob$Treatment == "Model"] <- "Predator Context"
bob$Test[bob$Treatment == "Mirror"] <- "Conspecific"
bob$Test[bob$Treatment == "Prey"] <- "Foraging Context"

Test_Order <- c('Control', 'Light Context', 'Foraging Context', 'Conspecific', 'Predator Context')

allstops <- subset(bobraw, bobraw$Behavior == "stopping")
all <- subset(allstops, allstops$Treatment == "Light"| allstops$Treatment =="Dark" |
                allstops$Treatment =="Model"  | allstops$Treatment =="Mirror"| allstops$Treatment =="Pre

all$Test[all$Treatment == "Light"] <- "Control"
all$Test[all$Treatment == "Dark"] <- "Light Context"
all$Test[all$Treatment == "Model"] <- "Predator Context"
all$Test[all$Treatment == "Mirror"] <- "Conspecific"
```

```
all$Test[all$Treatment == "Prey"] <- "Foraging Context"

all_filtered <- all[all$Duration <= 15, ]

allwalk <- subset(bobraw, bobraw$Behavior == "walking")
walk <- subset(allwalk, allwalk$Treatment == "Light"| allwalk$Treatment =="Dark" |
                allwalk$Treatment =="Model"  | allwalk$Treatment =="Mirror"| allwalk$Treatment =="Prey")

walk$Test[walk$Treatment == "Light"] <- "Control"
walk$Test[walk$Treatment == "Dark"] <- "Light Context"
walk$Test[walk$Treatment == "Model"] <- "Predator Context"
walk$Test[walk$Treatment == "Mirror"] <- "Conspecific"
walk$Test[walk$Treatment == "Prey"] <- "Foraging Context"

spiders <- subset(bobraw, bobraw$Treatment == "Light" | bobraw$Treatment =="Marpissa" | bobraw$Treatmen
```

```
spiderstops <- subset(spiders, spiders$Behavior == "stopping")

spiderstops$Treatment[spiderstops$Treatment == "Light"] <- "Heliophanus"
spiderstops$Treatment[spiderstops$Treatment == "Salticus"] <- "Salticus"
spiderstops$Treatment[spiderstops$Treatment == "Marpissa"] <- "Marpissa"

Spider_Order <- c('Heliophanus','Marpissa', 'Salticus')

filtered_data <- spiderstops[spiderstops$Duration <= 15, ]

bob$sex[bob$sex == "f"] <- "Female"
bob$sex[bob$sex == "m"] <- "Male"
bob$sex[bob$sex == "juv"] <- "Juvenile"

Sex_Order <- c('Juvenile', 'Female', 'Male')

bobcontrol <- subset(bob, bob$Test == "Control")
```

#### Gait description

#Since other spider species tested do not show any instances of abdomen bobbing, we compare the duration of gait stops between Heliophanus, Salticus and Marpissa (note: gait stops are all stops under 15 seconds)

#### Spiders Stopping Duration (Gait description)

```
ggplot(filtered_data, aes(factor(Treatment, level=Spider_Order), Duration, fill = Treatment))+
  geom_boxplot(outlier.shape = 1, outlier.size = 2)+
  geom_jitter(size = 0.8)+
  scale_fill_manual(values = cbbPalette)+
  xlab("Tested spider")+
  ylab("Duration of stops (s)")+
  ggtitle("Stop duration of tested spiders")+
  theme(plot.title = element_text(size = 15, face = "bold"))+
  ylim(0,16)
```

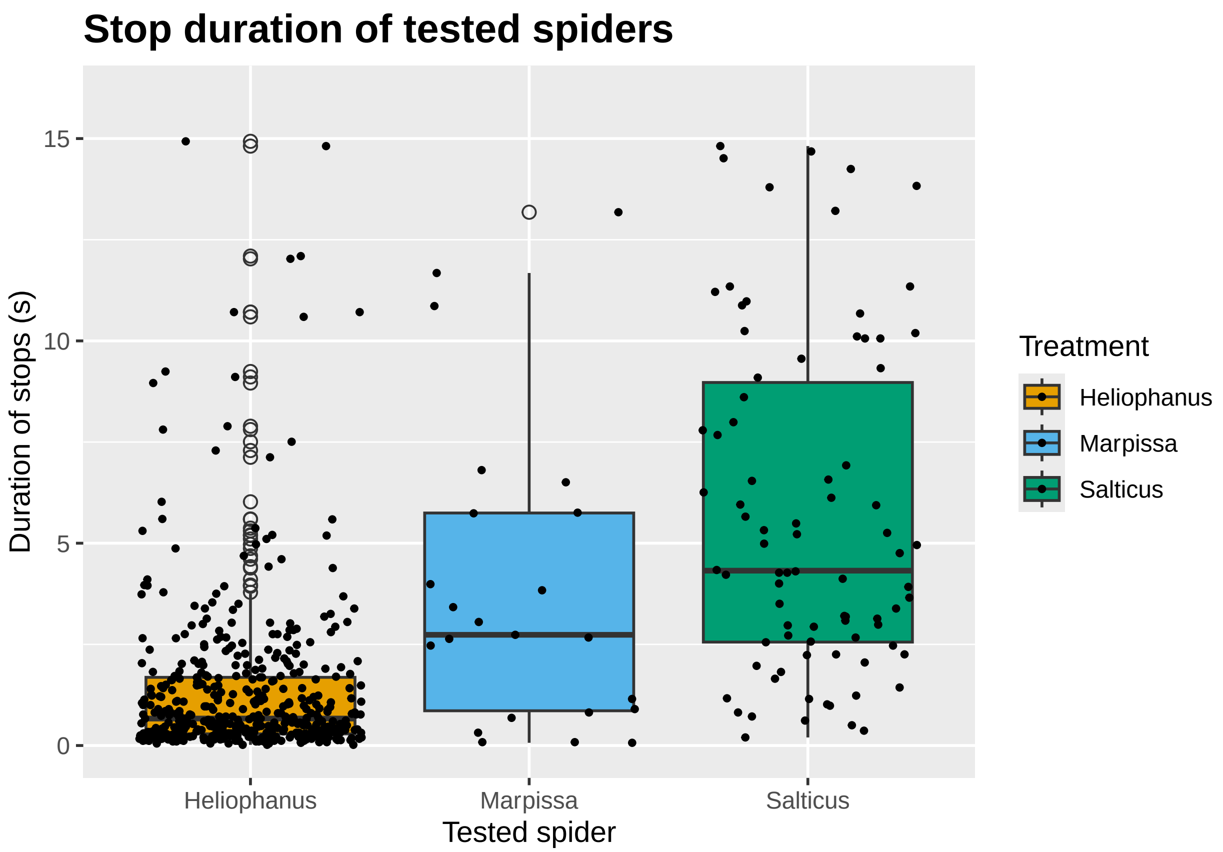


Gait stops seem to be longer in Salticus and Marpissa, let's see if significantly so by running a GLMM:

```
spiderstopmodel <- glmmTMB(Duration ~ Treatment + (1|Subject), data = filtered_data, family = Gamma(lin
simres <- simulateResiduals(spiderstopmodel)
plot(simres)
```

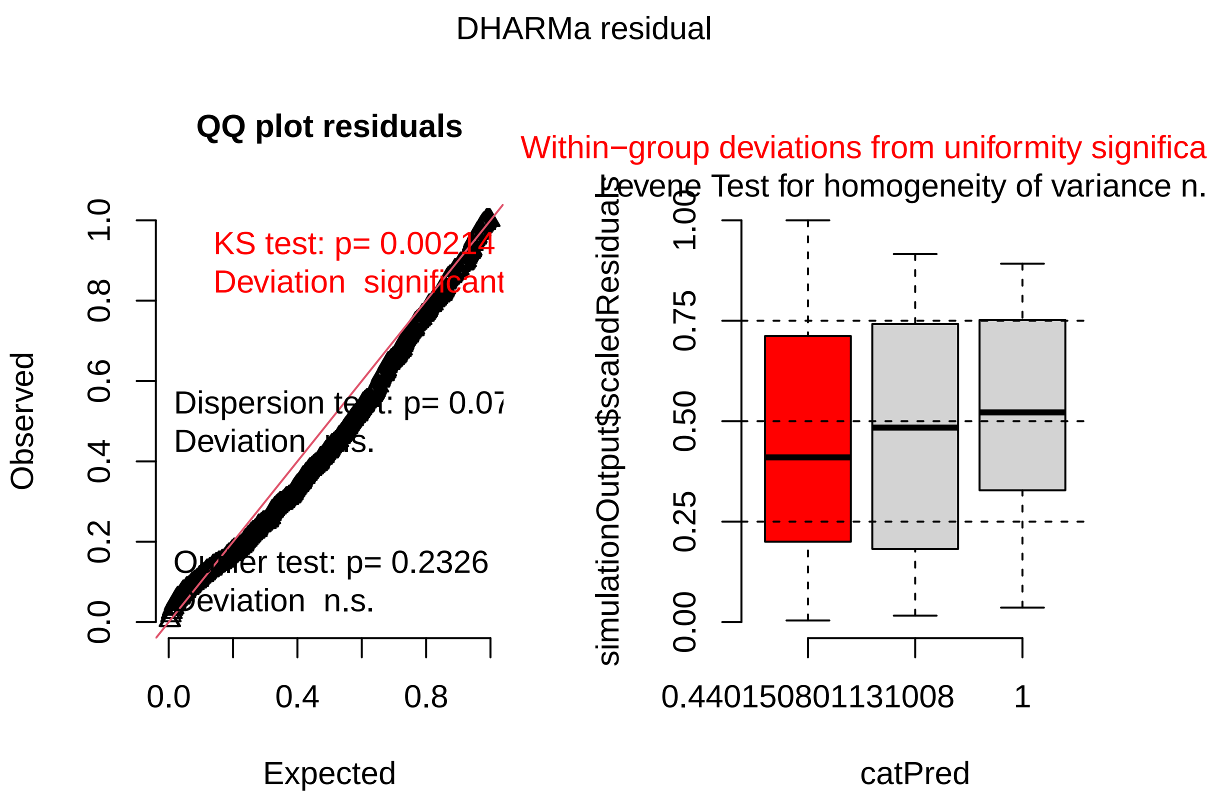


```r
Anova(spiderstopmodel)
```

```
## Analysis of Deviance Table (Type II Wald chisquare tests)
##
## Response: Duration
##            Chisq Df Pr(>Chisq)
## Treatment 83.479  2  < 2.2e-16 ***
## ---
## Signif. codes:  0 '***' 0.001 '**' 0.01 '*' 0.05 '.' 0.1 ' ' 1
```

```r
summary(spiderstopmodel)
```

```
##  Family: Gamma  ( log )
## Formula:          Duration ~ Treatment + (1 | Subject)
## Data: filtered_data
##
##      AIC       BIC    logLik -2*log(L)  df.resid
##   1747.7    1769.4    -868.8    1737.7       566
##
## Random effects:
##
## Conditional model:
##  Groups  Name        Variance Std.Dev.
##  Subject (Intercept) 0.1801   0.4244
```

```
## Number of obs: 571, groups:  Subject, 78
##
## Dispersion estimate for Gamma family (sigma^2): 0.942
##
## Conditional model:
##                    Estimate Std. Error z value Pr(>|z|)
## (Intercept)         0.25669    0.07645   3.358 0.000786 ***
## TreatmentMarpissa   1.20290    0.29104   4.133 3.58e-05 ***
## TreatmentSalticus   1.55289    0.18128   8.566  < 2e-16 ***
## ---
## Signif. codes:  0 '***' 0.001 '**' 0.01 '*' 0.05 '.' 0.1 ' ' 1
```

```
e1 <- emmeans(spiderstopmodel, ~Treatment, type = "response")
pairs(e1, adjust = "bonferroni")
```

```
##  contrast               ratio     SE  df null z.ratio p.value
##  Heliophanus / Marpissa 0.300 0.0874 Inf    1  -4.133  0.0001
##  Heliophanus / Salticus 0.212 0.0384 Inf    1  -8.566  <.0001
##  Marpissa / Salticus    0.705 0.2290 Inf    1  -1.077  0.8440
##
## P value adjustment: bonferroni method for 3 tests
## Tests are performed on the log scale
```

#### Visualisation and Models

#### Bobbing Behavior

Now, we visualize abdomen bobbing per stop (three random stops were picked for each test)

First, we look at whether bobbing is different between males, females and juveniles

```
sex_plot <- ggplot(bobcontrol, aes(factor(sex, level=Sex_Order), AB, fill = sex)) +
  geom_violin(trim = FALSE, alpha = 0.5, color = NA) +  # Violin shape with soft fill
  geom_boxplot(width = 0.3, outlier.shape = 1, outlier.size = 2, color = "black") +  # Thin boxplot on
  geom_jitter(width = 0.2, height = 0.05, size = 0.8)+
  coord_cartesian(ylim = c(0, NA))+
  theme_minimal() +
  scale_fill_manual(values = cbbPalette)+
  labs(title = "Abdomen bob by sex",
       y = "Abdomen bob per stop (three random stops)",
       x = "Sex") +
  theme(legend.position = "none")


sex_plot
```

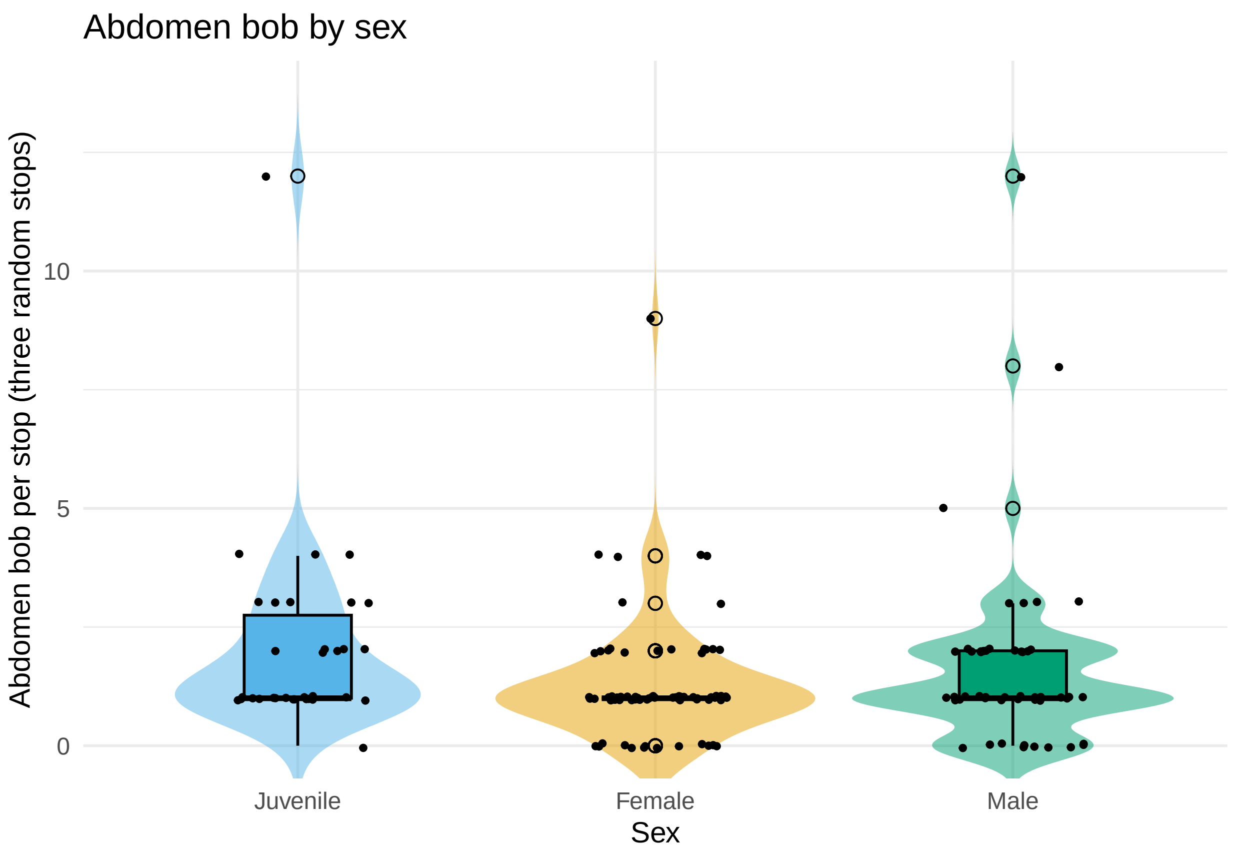


```
ggsave("sex_plot.svg", sex_plot, width = 8, height = 5, dpi = 300)
```

no difference can be detected between bobbing behavior in the control trials.

Let's run a model

```
sexmodel <- glmmTMB(AB ~ sex + (1|Subject), data = bobcontrol, family = poisson)
Anova(sexmodel)
```

```
## Analysis of Deviance Table (Type II Wald chisquare tests)
##
## Response: AB
##      Chisq Df Pr(>Chisq)
## sex 4.0234  2     0.1338
```

```
simres <- simulateResiduals(sexmodel)
plot(simres)
```

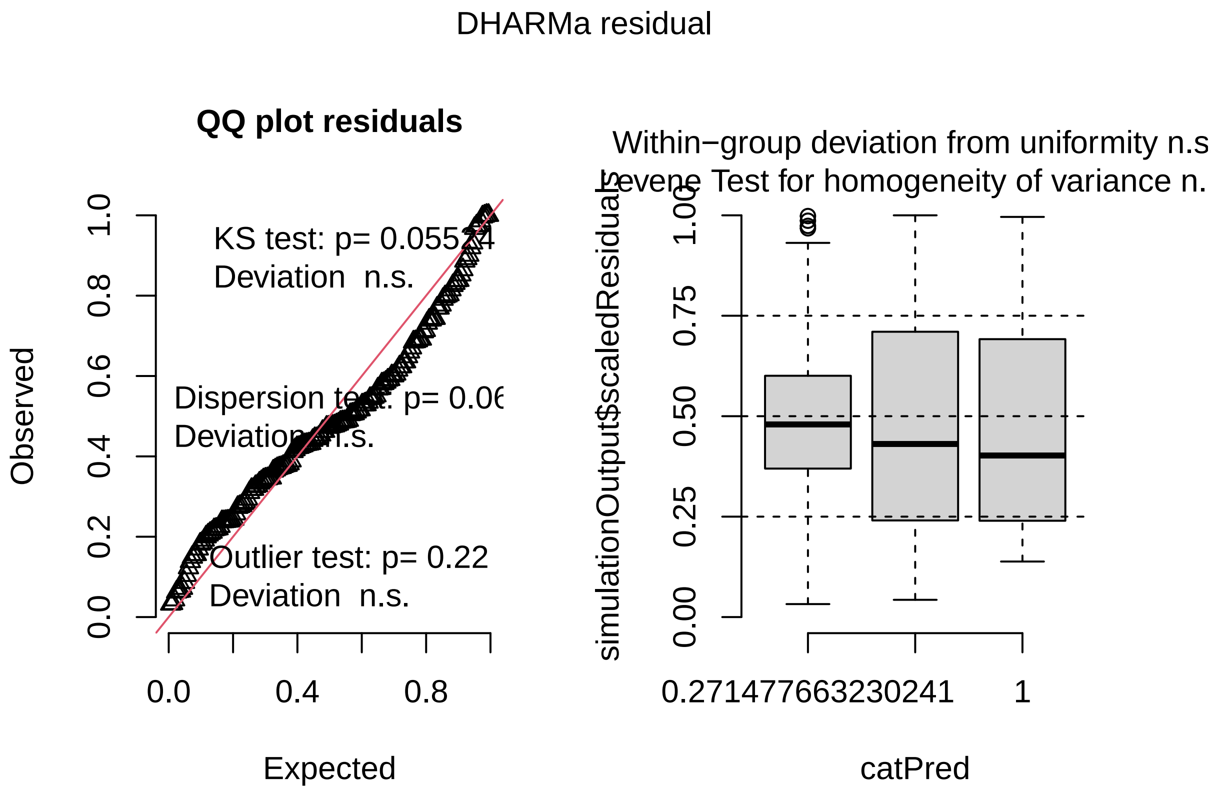


```r
summary(sexmodel)
```

```
##  Family: poisson  ( log )
## Formula:          AB ~ sex + (1 | Subject)
## Data: bobcontrol
##
##      AIC       BIC    logLik -2*log(L)  df.resid
##    526.6     539.0    -259.3     518.6       158
##
## Random effects:
##
## Conditional model:
##  Groups  Name        Variance Std.Dev.
##  Subject (Intercept) 0.1824   0.4271
## Number of obs: 162, groups:  Subject, 55
##
## Conditional model:
##             Estimate Std. Error z value Pr(>|z|)
## (Intercept)   0.1897     0.1323   1.434   0.1516
## sexJuvenile   0.4356     0.2187   1.992   0.0464 *
## sexMale       0.2000     0.2053   0.974   0.3299
## ---
## Signif. codes:  0 '***' 0.001 '**' 0.01 '*' 0.05 '.' 0.1 ' ' 1
```

```r
e5 <- emmeans(sexmodel, ~sex, type ="response")
pairs(e5)
```

```
##  contrast           ratio    SE  df null z.ratio p.value
##  Female / Juvenile 0.647 0.141 Inf    1  -1.992  0.1141
##  Female / Male     0.819 0.168 Inf    1  -0.974  0.5931
##  Juvenile / Male   1.266 0.299 Inf    1   0.997  0.5788
##
## P value adjustment: tukey method for comparing a family of 3 estimates
## Tests are performed on the log scale
```

There is no significant effect of bobbing behavior between males, females and juveniles.

Next, we look at bobbing behavior across the different treatments:

```r
bob_treatments <- ggplot(bob, aes(factor(Test, level = Test_Order), AB, fill = Treatment))+
  geom_boxplot(outlier.shape = 1, outlier.size = 2)+
  geom_jitter(size = 0.8)+
  scale_fill_manual(values = cbbPalette)+
  xlab("Treatment")+
  ylab("Abdomen bob per stop (three random stops)")+
  ggtitle("Abdomen bobbing per treatment")

bob_treatments
```

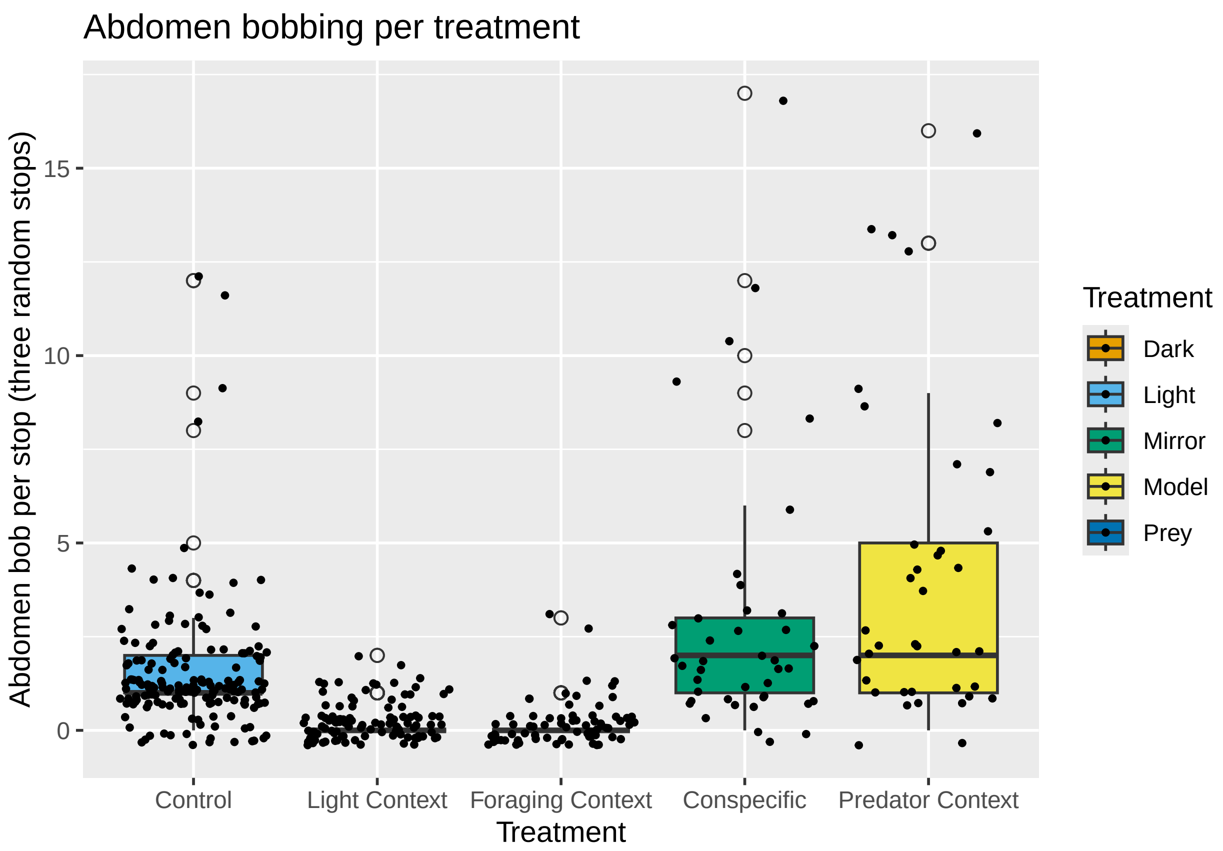

```r
ggsave("bob_treatments.svg", bob_treatments, width = 8, height = 5, dpi = 300)
```

We clearly see that there are very few abdomen bobs in the "dark" and the "prey" trial, we will now run a GLMM to further investigate this effect using a poisson distribution (as we are working with count data)

```r
bobmodel <- glmmTMB(AB ~ Test + (1|Subject), data = bob, family = poisson)
plot(simulateResiduals(bobmodel))
```

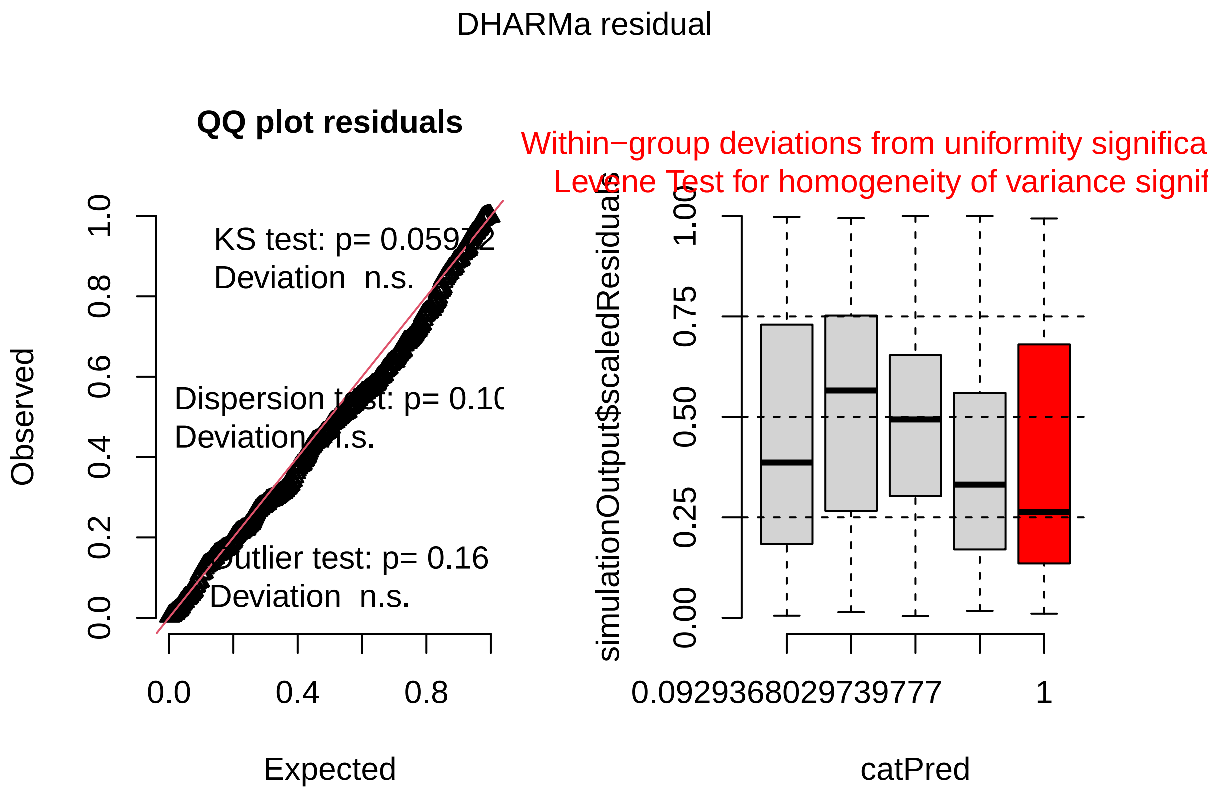


```r
Anova(bobmodel)
```

```
## Analysis of Deviance Table (Type II Wald chisquare tests)
##
## Response: AB
##       Chisq Df Pr(>Chisq)
## Test 184.2  4  < 2.2e-16 ***
## ---
## Signif. codes:  0 '***' 0.001 '**' 0.01 '*' 0.05 '.' 0.1 ' ' 1
```

```r
summary(bobmodel)
```

```
##  Family: poisson  ( log )
## Formula:          AB ~ Test + (1 | Subject)
## Data: bob
```

```
##
##       AIC       BIC    logLik -2*log(L)  df.resid
##    1196.9    1221.2    -592.4    1184.9       416
##
## Random effects:
##
## Conditional model:
##  Groups  Name        Variance Std.Dev.
##  Subject (Intercept) 0.2674   0.5171
## Number of obs: 422, groups:  Subject, 102
##
## Conditional model:
##                        Estimate Std. Error z value Pr(>|z|)
## (Intercept)              0.9753     0.1527   6.386 1.70e-10 ***
## TestControl             -0.6688     0.1632  -4.098 4.17e-05 ***
## TestForaging Context  -2.6645     0.3167  -8.414  < 2e-16 ***
## TestLight Context     -2.3959     0.2579  -9.290  < 2e-16 ***
## TestPredator Context   0.4192     0.1490   2.814   0.0049 **
## ---
## Signif. codes:  0 '***' 0.001 '**' 0.01 '*' 0.05 '.' 0.1 ' ' 1
```

While the distribution is not perfect, it looks ok, so we proceed to do a posthoc test using bonferroni correction

```
e2 <- emmeans(bobmodel, ~Test, type = "response")
pairs(e2, adjust = "bonferroni")
```

```
##  contrast                               ratio     SE  df null z.ratio p.value
##  Conspecific / Control                 1.9519 0.3190 Inf    1   4.098  0.0004
##  Conspecific / Foraging Context       14.3614 4.5500 Inf    1   8.414  <.0001
##  Conspecific / Light Context          10.9783 2.8300 Inf    1   9.290  <.0001
##  Conspecific / Predator Context        0.6576 0.0980 Inf    1  -2.814  0.0490
##  Control / Foraging Context            7.3575 2.1600 Inf    1   6.806  <.0001
##  Control / Light Context               5.6243 1.2400 Inf    1   7.833  <.0001
##  Control / Predator Context            0.3369 0.0532 Inf    1  -6.890  <.0001
##  Foraging Context / Light Context      0.7644 0.2700 Inf    1  -0.761  1.0000
##  Foraging Context / Predator Context   0.0458 0.0143 Inf    1  -9.904  <.0001
##  Light Context / Predator Context      0.0599 0.0151 Inf    1 -11.196  <.0001
##
## P value adjustment: bonferroni method for 10 tests
## Tests are performed on the log scale
```

We can clearly see that both dark and prey trials lie underneath the control baseline while bobbing increases during predator and conspecific encounters. The strongest effect is seen in trials with a model predator.

#### Stopping

random stops:

```
stop_treatment <- ggplot(bob, aes(factor(Test, level= Test_Order), Stop_Time, fill = Test))+
  geom_boxplot(outlier.shape = 1, outlier.size = 2)+
  geom_jitter(size = 0.8)+
```

```
  scale_fill_manual(values = cbbPalette)+
  xlab("Treatment")+
  ylab("Duration of stop (s)")+
  ggtitle("Stop duration per treatment")+
  theme(plot.title = element_text(size = 15, face = "bold"))

stop_treatment
```

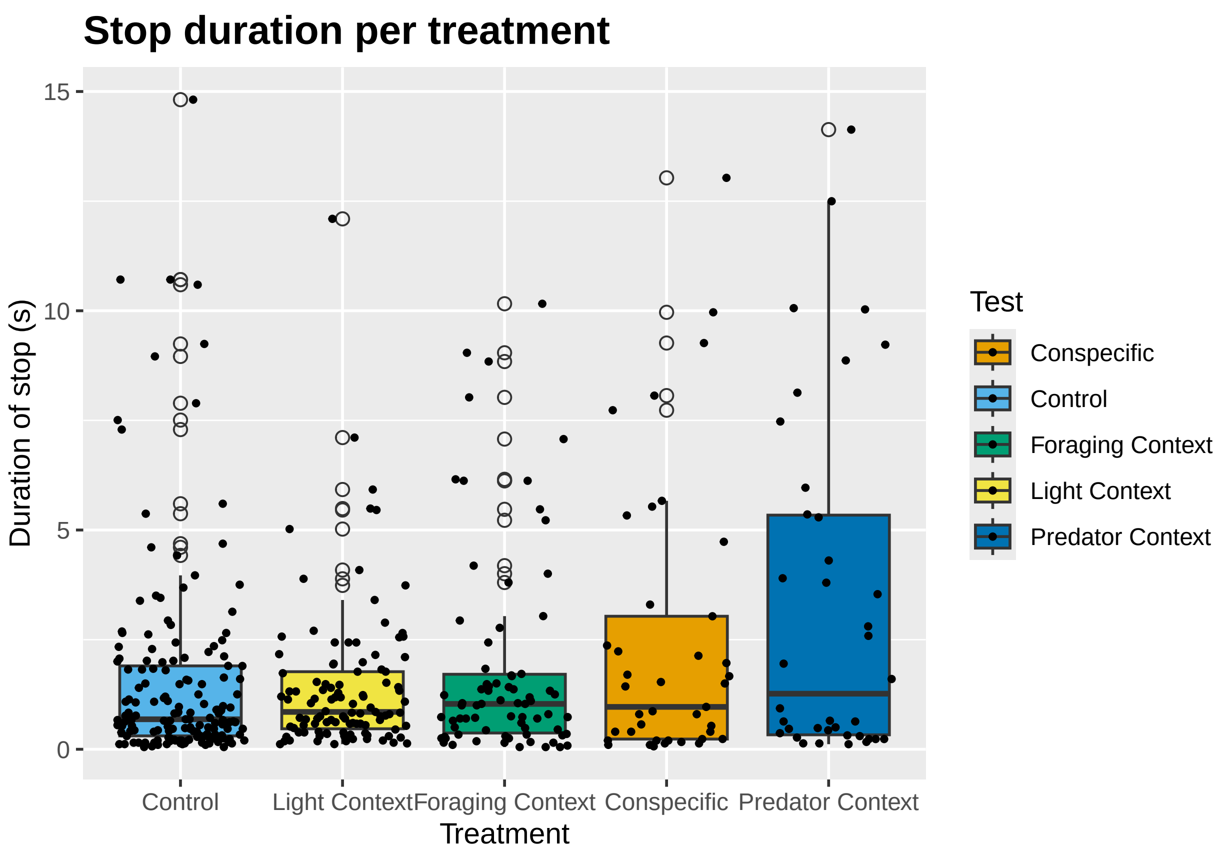


```
ggsave("stop_treatment.svg", stop_treatment, width = 8, height = 5, dpi = 300)
```

Here, we can see that the gait as such (i.e. stops) is quite consistent (especially the medians), with a higher variance in the predator and conspecific trials.

We will now run GLMM to investigate whether there are any significant effects.

```
stopmodel <- glmmTMB(Stop_Time ~ Test + (1|Subject), data = bob, family = Gamma(link = "log"))
plot(simulateResiduals(stopmodel))
```

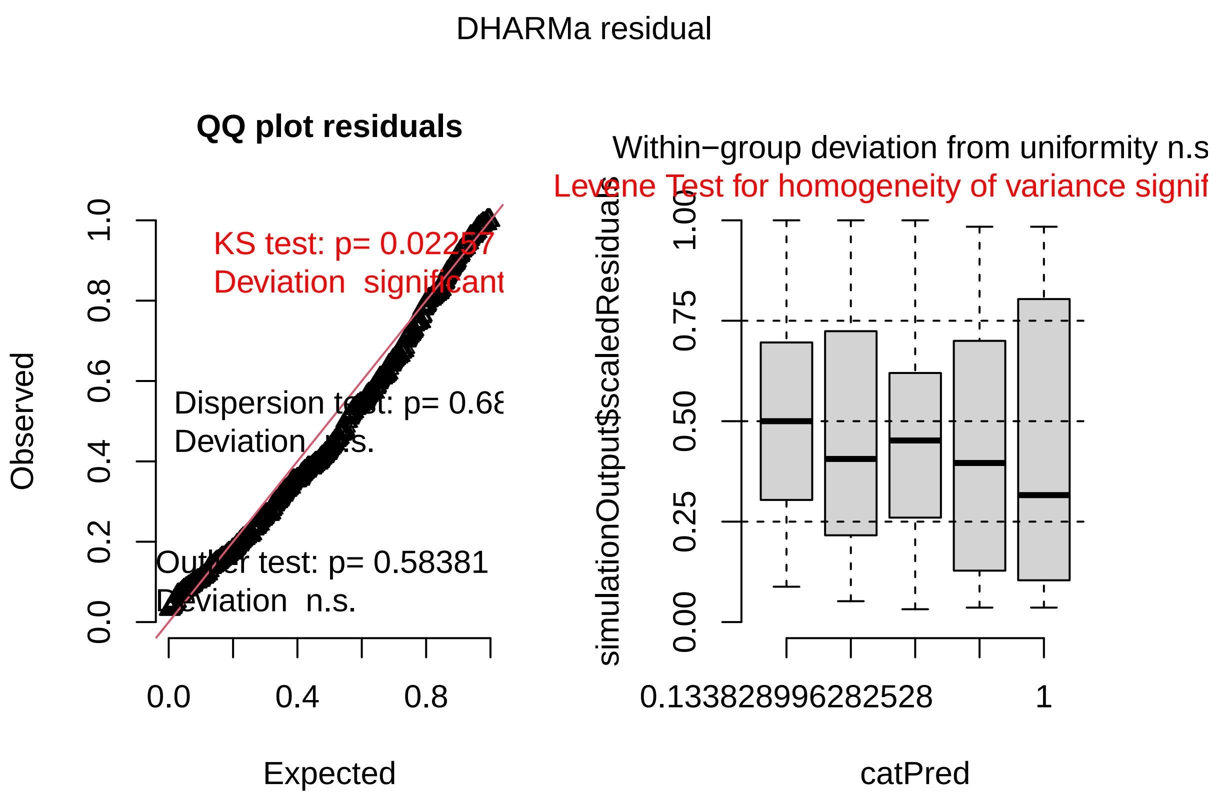


```r
Anova(stopmodel)
```

```
## Analysis of Deviance Table (Type II Wald chisquare tests)
##
## Response: Stop_Time
##       Chisq Df Pr(>Chisq)
## Test 19.094  4  0.0007533 ***
## ---
## Signif. codes:  0 '***' 0.001 '**' 0.01 '*' 0.05 '.' 0.1 ' ' 1
```

```r
summary(stopmodel)
```

```
##  Family: Gamma  ( log )
## Formula:          Stop_Time ~ Test + (1 | Subject)
## Data: bob
##
##      AIC      BIC   logLik -2*log(L)  df.resid
##   1325.5   1353.8   -655.7    1311.5       415
##
## Random effects:
##
## Conditional model:
##  Groups  Name        Variance Std.Dev.
##  Subject (Intercept) 0.1589   0.3986
```

```
## Number of obs: 422, groups:  Subject, 102
##
## Dispersion estimate for Gamma family (sigma^2): 1.08
##
## Conditional model:
##                        Estimate Std. Error z value Pr(>|z|)
## (Intercept)              0.9169     0.2054   4.464 8.04e-06 ***
## TestControl             -0.5467     0.2222  -2.461   0.0139 *
## TestForaging Context  -0.3659     0.2581  -1.418   0.1563
## TestLight Context      -0.5835     0.2384  -2.447   0.0144 *
## TestPredator Context   0.3109     0.2623   1.185   0.2358
## ---
## Signif. codes:  0 '***' 0.001 '**' 0.01 '*' 0.05 '.' 0.1 ' ' 1
```

While the distribution does not look perfect, it looks quite acceptable. There is a significant effect of Treatment on Stop duration, we will now investigate where the effect comes from using a posthoc test:

```
e3 <- emmeans(stopmodel, ~Test, type = "response")
pairs(e3, adjust = "bonferroni")
```

```
##  contrast                                ratio     SE  df null z.ratio p.value
##  Conspecific / Control                   1.727 0.3840 Inf    1   2.461  0.1387
##  Conspecific / Foraging Context          1.442 0.3720 Inf    1   1.418  1.0000
##  Conspecific / Light Context             1.792 0.4270 Inf    1   2.447  0.1439
##  Conspecific / Predator Context          0.733 0.1920 Inf    1  -1.185  1.0000
##  Control / Foraging Context              0.835 0.1520 Inf    1  -0.990  1.0000
##  Control / Light Context                 1.038 0.1560 Inf    1   0.245  1.0000
##  Control / Predator Context              0.424 0.0942 Inf    1  -3.861  0.0011
##  Foraging Context / Light Context        1.243 0.2460 Inf    1   1.098  1.0000
##  Foraging Context / Predator Context 0.508 0.1290 Inf    1  -2.671  0.0757
##  Light Context / Predator Context    0.409 0.0958 Inf    1  -3.816  0.0014
##
## P value adjustment: bonferroni method for 10 tests
## Tests are performed on the log scale
```

#### All stops (raw data)

Now, we will investigate whether the duration of stops changes with different treatments, i.e. whether the gait changes. First, we plot the raw data (now, we include all recorded stops under 15 seconds, not only three random stops)

```
allstops <- ggplot(all, aes(factor(Test, level = Test_Order), Duration, fill = Test))+
  geom_boxplot()+
  geom_jitter(size = 0.8)+
  scale_fill_manual(values = cbbPalette)+
  xlab("Treatment")+
  ylab("Duration of stop (s)")+
  ggtitle("Stop duration across all treatment")+
  theme(plot.title = element_text(size = 15, face = "bold"))+
  ylim(0,20)

allstops
```

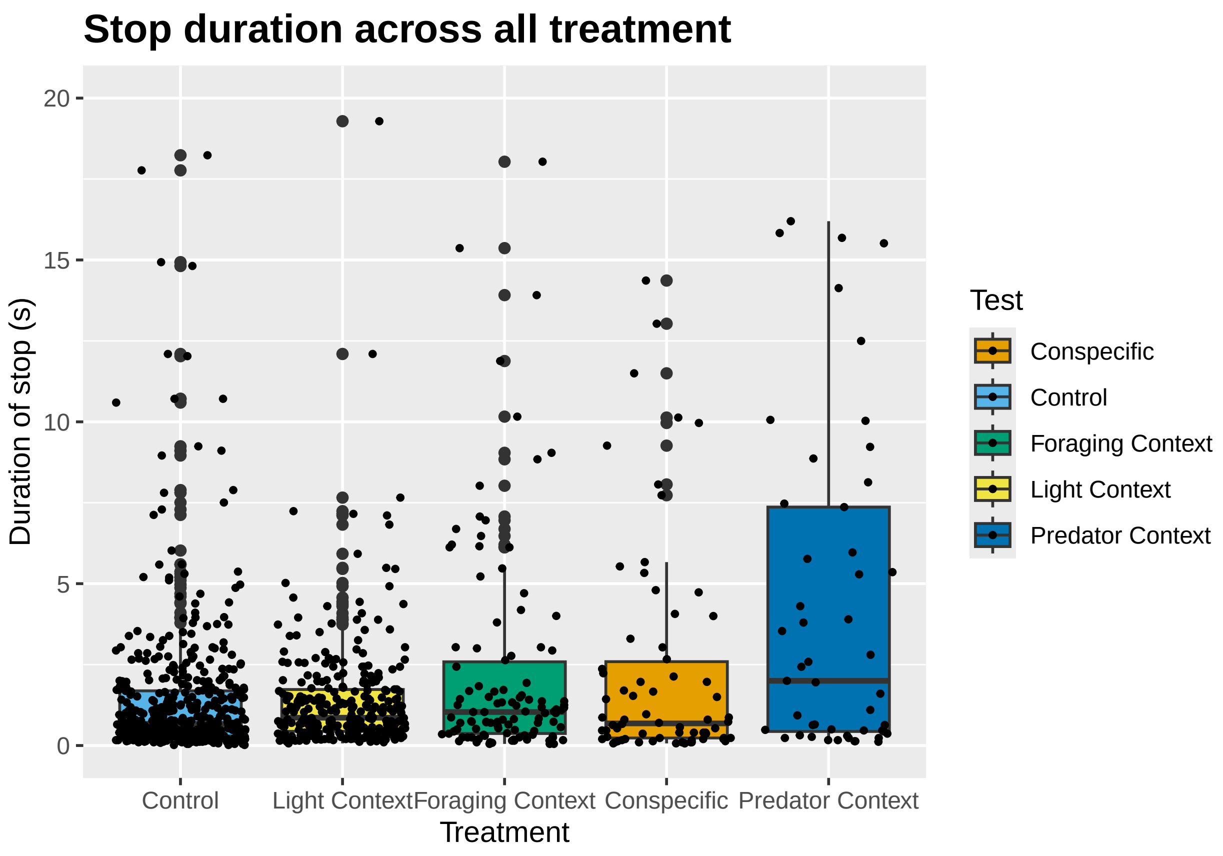


```
ggsave("all_stops.svg", allstops, width = 8, height = 5, dpi = 300)
```

```
# Filter for Treatment == "Light"
light_data <- bobraw %>% filter(Treatment == "Light")

# Separate behaviors
abdomen_behaviors <- light_data %>%
  filter(Behavior %in% c("abdomen down", "abdomen up"))

stopping_behaviors <- light_data %>%
  filter(Behavior == "stopping", Duration < 15)

# Check for overlap per stopping row
stopping_with_overlap <- stopping_behaviors %>%
  rowwise() %>%
  mutate(overlap = any(
    abdomen_behaviors$Subject == Subject &
    abdomen_behaviors$Image.index.start <= Image.index.stop &
    abdomen_behaviors$Image.index.stop >= Image.index.start
  )) %>%
  ungroup()

# Calculate percentage
percent_stops_with_abdomen <- mean(stopping_with_overlap$overlap) * 100
```

```r
walkcont <- subset(walk, walk$Treatment == "Light")
describe.by(walkcont, walkcont$Behavior)
```

```
##
##  Descriptive statistics by group
## group: walking
##                    vars   n    mean      sd  median trimmed     mad    min
## Observation.id        1 539   29.83   15.48   32.00   30.40   19.27   1.00
## Subject               2 539   30.23   17.05   33.00   30.59   22.24   1.00
## Behavior              3 539    1.00    0.00    1.00    1.00    0.00   1.00
## Behavior.type         4 539    1.00    0.00    1.00    1.00    0.00   1.00
## Start                 5 539   79.64   52.52   67.82   71.46   43.31  15.25
## Stop                  6 539   81.92   52.98   69.77   73.66   42.94  16.73
## Duration              7 539    2.28    3.75    1.08    1.46    1.06   0.02
## Image.index.start     8 539 4688.84 3190.40 3984.00 4188.08 2661.27 616.00
## Image.index.stop      9 539 4824.39 3220.99 4106.00 4322.02 2639.03 619.00
## Treatment            10 539    1.00    0.00    1.00    1.00    0.00   1.00
## Trial                11 539    1.48    0.50    1.00    1.47    0.00   1.00
## Date                 12 539    6.54    2.59    8.00    6.58    1.48   1.00
## Test                 13 539    1.00    0.00    1.00    1.00    0.00   1.00
##                         max    range  skew kurtosis     se
## Observation.id        56.00    55.00 -0.27    -1.12   0.67
## Subject               55.00    54.00 -0.16    -1.40   0.73
## Behavior               1.00     0.00   NaN      NaN   0.00
## Behavior.type          1.00     0.00   NaN      NaN   0.00
## Start                290.27   275.02  1.49     2.33   2.26
## Stop                 300.95   284.22  1.49     2.32   2.28
## Duration              33.07    33.05  4.40    25.04   0.16
## Image.index.start 17399.00 16783.00  1.48     2.25 137.42
## Image.index.stop  18039.00 17420.00  1.47     2.23 138.74
## Treatment              1.00     0.00   NaN      NaN   0.00
## Trial                  2.00     1.00  0.09    -2.00   0.02
## Date                  12.00    11.00 -0.36    -0.87   0.11
## Test                   1.00     0.00   NaN      NaN   0.00
```

```r
describe(walkcont, IQR = TRUE)
```

```
##                    vars   n    mean      sd  median trimmed     mad    min
## Observation.id*       1 539   29.83   15.48   32.00   30.40   19.27   1.00
## Subject*              2 539   30.23   17.05   33.00   30.59   22.24   1.00
## Behavior*             3 539    1.00    0.00    1.00    1.00    0.00   1.00
## Behavior.type*        4 539    1.00    0.00    1.00    1.00    0.00   1.00
## Start                 5 539   79.64   52.52   67.82   71.46   43.31  15.25
## Stop                  6 539   81.92   52.98   69.77   73.66   42.94  16.73
## Duration              7 539    2.28    3.75    1.08    1.46    1.06   0.02
## Image.index.start     8 539 4688.84 3190.40 3984.00 4188.08 2661.27 616.00
## Image.index.stop      9 539 4824.39 3220.99 4106.00 4322.02 2639.03 619.00
## Treatment*           10 539    1.00    0.00    1.00    1.00    0.00   1.00
## Trial                11 539    1.48    0.50    1.00    1.47    0.00   1.00
## Date*                12 539    6.54    2.59    8.00    6.58    1.48   1.00
## Test*                13 539    1.00    0.00    1.00    1.00    0.00   1.00
##                         max    range  skew kurtosis     se     IQR
```

```
## Observation.id*      56.00    55.00 -0.27    -1.12   0.67   25.50
## Subject*             55.00    54.00 -0.16    -1.40   0.73   33.00
## Behavior*             1.00     0.00   NaN      NaN   0.00    0.00
## Behavior.type*        1.00     0.00   NaN      NaN   0.00    0.00
## Start               290.27   275.02  1.49     2.33   2.26   60.45
## Stop                300.95   284.22  1.49     2.32   2.28   59.07
## Duration             33.07    33.05  4.40    25.04   0.16    1.87
## Image.index.start 17399.00 16783.00  1.48     2.25 137.42 3687.00
## Image.index.stop  18039.00 17420.00  1.47     2.23 138.74 3661.00
## Treatment*            1.00     0.00   NaN      NaN   0.00    0.00
## Trial                 2.00     1.00  0.09    -2.00   0.02    1.00
## Date*                12.00    11.00 -0.36    -0.87   0.11    3.00
## Test*                 1.00     0.00   NaN      NaN   0.00    0.00
```

```r
stopcont <- subset(all_filtered, all$Treatment == "Light")
describe.by(stopcont, stopcont$Behavior)
```

```
##
##  Descriptive statistics by group
## group: stopping
##                   vars   n    mean      sd  median trimmed     mad     min
## Observation.id       1 466   31.42   17.29   34.00   31.56   20.76    1.00
## Subject              2 466   31.82   18.26   34.00   32.20   25.20    1.00
## Behavior             3 466    1.00    0.00    1.00    1.00    0.00    1.00
## Behavior.type        4 466    1.00    0.00    1.00    1.00    0.00    1.00
## Start                5 466   82.75   52.01   71.26   74.72   42.87   16.73
## Stop                 6 466   84.30   52.11   72.91   76.24   42.36   17.08
## Duration             7 466    1.55    2.27    0.67    1.03    0.69    0.02
## Image.index.start    8 466 4857.55 3086.31 4112.00 4366.61 2338.06 1003.00
## Image.index.stop     9 466 4947.83 3089.19 4189.00 4455.66 2336.58 1024.00
## Treatment           10 466    2.11    0.44    2.00    2.00    0.00    1.00
## Trial               11 466    1.46    0.50    1.00    1.45    0.00    1.00
## Date                12 466    7.16    3.18    9.00    7.16    1.48    1.00
## Test                13 466    1.14    0.56    1.00    1.00    0.00    1.00
##                        max    range  skew kurtosis     se
## Observation.id       64.00    63.00 -0.12    -1.09   0.80
## Subject              58.00    57.00 -0.16    -1.41   0.85
## Behavior              1.00     0.00   NaN      NaN   0.00
## Behavior.type         1.00     0.00   NaN      NaN   0.00
## Start               289.74   273.01  1.44     2.06   2.41
## Stop                290.27   273.19  1.45     2.09   2.41
## Duration             14.93    14.92  3.09    11.14   0.11
## Image.index.start 17367.00 16364.00  1.52     2.34 142.97
## Image.index.stop  17399.00 16375.00  1.53     2.36 143.10
## Treatment             4.00     3.00  3.41    11.97   0.02
## Trial                 2.00     1.00  0.15    -1.98   0.02
## Date                 16.00    15.00 -0.16    -0.78   0.15
## Test                  4.00     3.00  4.26    17.85   0.03
```

```r
describe(stopcont, IQR = TRUE)
```

```
##                   vars   n    mean      sd  median trimmed     mad     min
## Observation.id*      1 466   31.42   17.29   34.00   31.56   20.76    1.00
```

```
## Subject*               2 466   31.82   18.26   34.00   32.20   25.20    1.00
## Behavior*              3 466    1.00    0.00    1.00    1.00    0.00    1.00
## Behavior.type*         4 466    1.00    0.00    1.00    1.00    0.00    1.00
## Start                  5 466   82.75   52.01   71.26   74.72   42.87   16.73
## Stop                   6 466   84.30   52.11   72.91   76.24   42.36   17.08
## Duration               7 466    1.55    2.27    0.67    1.03    0.69    0.02
## Image.index.start      8 466 4857.55 3086.31 4112.00 4366.61 2338.06 1003.00
## Image.index.stop       9 466 4947.83 3089.19 4189.00 4455.66 2336.58 1024.00
## Treatment*            10 466    2.11    0.44    2.00    2.00    0.00    1.00
## Trial                 11 466    1.46    0.50    1.00    1.45    0.00    1.00
## Date*                 12 466    7.16    3.18    9.00    7.16    1.48    1.00
## Test*                 13 466    1.14    0.56    1.00    1.00    0.00    1.00
##                         max    range  skew kurtosis     se     IQR
## Observation.id*       64.00    63.00 -0.12    -1.09   0.80   28.00
## Subject*              58.00    57.00 -0.16    -1.41   0.85   36.00
## Behavior*              1.00     0.00   NaN      NaN   0.00    0.00
## Behavior.type*         1.00     0.00   NaN      NaN   0.00    0.00
## Start                289.74   273.01  1.44     2.06   2.41   59.05
## Stop                 290.27   273.19  1.45     2.09   2.41   57.70
## Duration              14.93    14.92  3.09    11.14   0.11    1.51
## Image.index.start 17367.00 16364.00  1.52     2.34 142.97 3320.00
## Image.index.stop  17399.00 16375.00  1.53     2.36 143.10 3244.25
## Treatment*             4.00     3.00  3.41    11.97   0.02    0.00
## Trial                  2.00     1.00  0.15    -1.98   0.02    1.00
## Date*                 16.00    15.00 -0.16    -0.78   0.15    4.00
## Test*                  4.00     3.00  4.26    17.85   0.03    0.00
```

```r
describe(filtered_data, IQR = TRUE)
```

```
##                   vars   n    mean      sd  median trimmed     mad    min
## Observation.id*      1 571   43.51   24.62   49.00   44.41   31.13   1.00
## Subject*             2 571   36.58   21.36   39.00   36.21   23.72   1.00
## Behavior*            3 571    1.00    0.00    1.00    1.00    0.00   1.00
## Behavior.type*       4 571    1.00    0.00    1.00    1.00    0.00   1.00
## Start                5 571   88.02   53.82   74.72   81.72   51.00  11.99
## Stop                 6 571   90.12   54.43   75.54   83.69   51.30  13.81
## Duration             7 571    2.10    2.93    0.90    1.41    1.04   0.02
## Image.index.start    8 571 5206.79 3270.19 4446.00 4825.40 3205.38 619.00
## Image.index.stop     9 571 5331.39 3309.07 4486.00 4943.68 3221.69 631.00
## Treatment*          10 571    1.33    0.71    1.00    1.16    0.00   1.00
## Trial               11 571    1.42    0.49    1.00    1.39    0.00   1.00
## Date*               12 571    9.12    3.65    9.00    9.28    4.45   1.00
##                         max    range  skew kurtosis     se     IQR
## Observation.id*       81.00    80.00 -0.25    -1.37   1.03   46.00
## Subject*              78.00    77.00  0.01    -1.14   0.89   38.00
## Behavior*              1.00     0.00   NaN      NaN   0.00    0.00
## Behavior.type*         1.00     0.00   NaN      NaN   0.00    0.00
## Start                289.74   277.74  1.02     0.70   2.25   72.57
## Stop                 290.27   276.46  1.02     0.62   2.28   73.24
## Duration              14.93    14.92  2.38     5.60   0.12    2.22
## Image.index.start 17367.00 16748.00  1.01     0.63 136.85 4547.50
## Image.index.stop  17399.00 16768.00  1.00     0.55 138.48 4553.50
## Treatment*             3.00     2.00  1.81     1.42   0.03    0.00
## Trial                  2.00     1.00  0.34    -1.89   0.02    1.00
```

```
## Date*                 16.00    15.00 -0.34    -1.02   0.15    7.00
```

```
spiderstops %>%
  group_by(Treatment) %>%
  summarise(n_subjects = n_distinct(Subject))
```

```
## # A tibble: 3 x 2
##   Treatment   n_subjects
##   <chr>            <int>
## 1 Heliophanus         55
## 2 Marpissa            10
## 3 Salticus            17
```

```
describe.by(bobcontrol, bobcontrol$Treatment)
```

```
##
##  Descriptive statistics by group
## group: Light
##             vars   n  mean    sd median trimmed   mad  min   max range  skew
## ID             1 162 28.81 16.07  29.00   28.92 20.76 1.00 56.00 55.00 -0.04
## Subject        2 162 28.49 15.81  29.00   28.58 20.76 1.00 55.00 54.00 -0.05
## Treatment      3 162  1.00  0.00   1.00    1.00  0.00 1.00  1.00  0.00   NaN
## Stop_Time      4 162  1.59  2.34   0.68    1.05  0.72 0.05 14.81 14.76  2.95
## AB             5 162  1.56  1.73   1.00    1.28  0.00 0.00 12.00 12.00  3.61
## AB_per_stop    6 162  2.32  2.43   1.53    1.90  1.70 0.00 14.93 14.93  1.89
## Date           7 162  7.62  4.01   7.00    7.77  5.93 1.00 12.00 11.00 -0.11
## Trial          8 162  1.42  0.50   1.00    1.40  0.00 1.00  2.00  1.00  0.32
## sex            9 162  1.83  0.87   2.00    1.78  1.48 1.00  3.00  2.00  0.34
## Test          10 162  1.00  0.00   1.00    1.00  0.00 1.00  1.00  0.00   NaN
##             kurtosis   se
## ID             -1.20 1.26
## Subject        -1.21 1.24
## Treatment        NaN 0.00
## Stop_Time       9.79 0.18
## AB             17.06 0.14
## AB_per_stop     4.71 0.19
## Date           -1.64 0.31
## Trial          -1.91 0.04
## sex            -1.62 0.07
## Test             NaN 0.00
```

```
allstops <- subset(bobraw, bobraw$Behavior == "stopping")
all <- subset(allstops, allstops$Treatment == "Light" | allstops$Treatment == "Salticus"| allstops$Trea
all_filtered <- all[all$Duration <= 15, ]

describe(all_filtered, IQR = TRUE)
```

```
##                   vars   n    mean      sd  median trimmed     mad    min
## Observation.id*      1 571   43.51   24.62   49.00   44.41   31.13   1.00
## Subject*             2 571   36.58   21.36   39.00   36.21   23.72   1.00
## Behavior*            3 571    1.00    0.00    1.00    1.00    0.00   1.00
## Behavior.type*       4 571    1.00    0.00    1.00    1.00    0.00   1.00
```

```
## Start                5 571   88.02   53.82   74.72   81.72   51.00  11.99
## Stop                 6 571   90.12   54.43   75.54   83.69   51.30  13.81
## Duration             7 571    2.10    2.93    0.90    1.41    1.04   0.02
## Image.index.start    8 571 5206.79 3270.19 4446.00 4825.40 3205.38 619.00
## Image.index.stop     9 571 5331.39 3309.07 4486.00 4943.68 3221.69 631.00
## Treatment*          10 571    1.33    0.71    1.00    1.16    0.00   1.00
## Trial               11 571    1.42    0.49    1.00    1.39    0.00   1.00
## Date*               12 571    9.12    3.65    9.00    9.28    4.45   1.00
##                        max    range  skew kurtosis     se     IQR
## Observation.id*      81.00    80.00 -0.25    -1.37   1.03   46.00
## Subject*             78.00    77.00  0.01    -1.14   0.89   38.00
## Behavior*             1.00     0.00   NaN      NaN   0.00    0.00
## Behavior.type*        1.00     0.00   NaN      NaN   0.00    0.00
## Start               289.74   277.74  1.02     0.70   2.25   72.57
## Stop                290.27   276.46  1.02     0.62   2.28   73.24
## Duration             14.93    14.92  2.38     5.60   0.12    2.22
## Image.index.start 17367.00 16748.00  1.01     0.63 136.85 4547.50
## Image.index.stop  17399.00 16768.00  1.00     0.55 138.48 4553.50
## Treatment*            3.00     2.00  1.81     1.42   0.03    0.00
## Trial                 2.00     1.00  0.34    -1.89   0.02    1.00
## Date*                16.00    15.00 -0.34    -1.02   0.15    7.00
```

```
salt <- subset(all_filtered, all_filtered$Treatment == "Salticus")
marp <- subset(all_filtered, all_filtered$Treatment == "Marpissa")

describe(salt, IQR = TRUE)
```

```
##                   vars  n    mean      sd  median trimmed     mad    min
## Observation.id*      1 82    9.82    4.52    8.00    9.82    2.97   1.00
## Subject*             2 82    8.54    4.12    9.00    8.42    3.71   1.00
## Behavior*            3 82    1.00    0.00    1.00    1.00    0.00   1.00
## Behavior.type*       4 82    1.00    0.00    1.00    1.00    0.00   1.00
## Start                5 82  129.99   46.28  138.29  133.76   36.90  11.99
## Stop                 6 82  135.66   47.05  142.66  138.92   39.40  13.81
## Duration             7 82    5.67    4.08    4.32    5.29    3.43   0.20
## Image.index.start    8 82 7791.73 2773.97 8289.00 8017.77 2212.04 719.00
## Image.index.stop     9 82 8131.72 2819.97 8551.00 8326.76 2361.78 828.00
## Treatment*          10 82    1.00    0.00    1.00    1.00    0.00   1.00
## Trial               11 82    1.07    0.26    1.00    1.00    0.00   1.00
## Date*               12 82    2.27    0.63    2.00    2.33    0.00   1.00
##                        max    range  skew kurtosis     se     IQR
## Observation.id*      18.00    17.00  0.15    -1.03   0.50    7.00
## Subject*             17.00    16.00  0.12    -0.80   0.45    6.75
## Behavior*             1.00     0.00   NaN      NaN   0.00    0.00
## Behavior.type*        1.00     0.00   NaN      NaN   0.00    0.00
## Start               217.07   205.07 -0.72     0.23   5.11   49.56
## Stop                227.31   213.50 -0.62     0.11   5.20   50.68
## Duration             14.81    14.62  0.70    -0.64   0.45    6.42
## Image.index.start 13011.00 12292.00 -0.72     0.23 306.33 2970.75
## Image.index.stop  13625.00 12797.00 -0.62     0.11 311.41 3037.50
## Treatment*            1.00     0.00   NaN      NaN   0.00    0.00
## Trial                 2.00     1.00  3.22     8.46   0.03    0.00
## Date*                 3.00     2.00 -0.26    -0.71   0.07    1.00
```

```r
describe(marp, IQR = TRUE)
```

```
##                     vars  n    mean      sd  median trimmed     mad     min
## Observation.id*        1 23    3.13    1.94    3.00    3.00    2.97    1.00
## Subject*               2 23    3.04    1.30    3.00    3.00    1.48    1.00
## Behavior*              3 23    1.00    0.00    1.00    1.00    0.00    1.00
## Behavior.type*         4 23    1.00    0.00    1.00    1.00    0.00    1.00
## Start                  5 23  119.03   60.94  122.06  116.11   74.43   34.52
## Stop                   6 23  122.92   59.63  123.21  119.90   74.60   37.94
## Duration               7 23    3.89    3.80    2.74    3.39    3.04    0.07
## Image.index.start      8 23 7134.87 3652.80 7316.00 6959.47 4461.14 2069.00
## Image.index.stop       9 23 7367.96 3574.51 7385.00 7186.79 4471.52 2274.00
## Treatment*            10 23    1.00    0.00    1.00    1.00    0.00    1.00
## Trial                 11 23    1.70    0.47    2.00    1.74    0.00    1.00
## Date*                 12 23    1.70    0.47    2.00    1.74    0.00    1.00
##                          max    range  skew kurtosis     se     IQR
## Observation.id*         7.00     6.00  0.29    -1.41   0.40    4.00
## Subject*                6.00     5.00  0.04    -0.41   0.27    1.50
## Behavior*               1.00     0.00   NaN      NaN   0.00    0.00
## Behavior.type*          1.00     0.00   NaN      NaN   0.00    0.00
## Start                 266.03   231.51  0.32    -0.65  12.71  105.02
## Stop                  268.50   230.56  0.37    -0.58  12.43  101.40
## Duration               13.18    13.11  1.06     0.08   0.79    4.89
## Image.index.start   15946.00 13877.00  0.32    -0.65 761.66 6295.00
## Image.index.stop    16094.00 13820.00  0.37    -0.58 745.34 6078.00
## Treatment*              1.00     0.00   NaN      NaN   0.00    0.00
## Trial                   2.00     1.00 -0.80    -1.42   0.10    1.00
## Date*                   2.00     1.00 -0.80    -1.42   0.10    1.00
```
